# Between heuristics and optimality: Flexible integration of cost and evidence during information sampling

**DOI:** 10.1101/2022.05.17.492355

**Authors:** Abigail Hsiung, John M. Pearson, Jia-Hou Poh, Shabnam Hakimi, R. Alison Adcock, Scott A. Huettel

## Abstract

Effective decision making in an uncertain world requires balancing the benefits of acquiring relevant information with the costs of delaying choice. Optimal strategies for information sampling can be accurate but computationally expensive, whereas heuristic strategies are often computationally simple but rigid. To characterize the computations that underlie information sampling, we examined choice processes in human participants who sampled sequences of images (e.g. indoor and outdoor scenes) and attempted to infer the majority category (e.g. indoor or outdoor) under two reward conditions. We examined how behavior maps onto potential information sampling strategies. We found that choices were best described by a flexible function that lay between optimality and heuristics; integrating the magnitude of evidence favoring each category and the number of samples collected thus far. Integration of these criteria resulted in a trade-off between evidence and samples collected, in which the strength of evidence needed to stop sampling decreased linearly as the number of samples accumulated over the course of a trial. This non-optimal trade-off best accounted for choice behavior even under high reward contexts. Our results demonstrate that unlike the optimal strategy, humans are performing simple accumulations instead of computing expected values, and that unlike a simple heuristic strategy, humans are dynamically integrating multiple sources of information in lieu of using only one source. This evidence-by-costs tradeoff illustrates a computationally efficient strategy that balances competing motivations for accuracy and cost minimization.

## Introduction

Before making important decisions, humans often collect information about the likely outcomes of different choice options. Consider the choice between two popular restaurants in a new city. Collecting information about both restaurants can increase the likelihood of a positive dining experience but also carries costs (e.g., time spent on evidence accumulation increases the likelihood that the options will become unavailable). Effective decision making thus requires information sampling strategies that balance accuracy and sampling costs – and understanding this balance remains a critical topic for decision science (Averbeck, 2015; Blanchard & Gershman, 2018; Cohen et al., 2007; Gigerenzer & Goldstein, 1996; Gold & Shadlen, 2007; Kolling et al., 2012; H. A. Simon, 1990). In particular, how is information about evidence and costs transformed into the decision to *sample* information or *stop*?

Current normative models of sequential information sampling posit that an optimal information sampling strategy should compare the expected values of available actions (continue sampling, choose option A, or choose option B) before selecting the action with the highest expected value (Coenen & Gureckis, 2016; Furl & Averbeck, 2011; Hauser et al., 2017, 2018; Moutoussis et al., 2011). Computing these expected values requires several steps. The decision maker must first determine the expected value of stopping by calculating the probability that each option is correct given the available evidence collected thus far. This estimate must, then, be multiplied by the available reward minus the costs accrued. To determine the value of continuing, the decision maker must estimate the expected values of potential future states, as if an additional sample was drawn. Accurate estimation of these future expected values necessitates an extensive backward induction process (Bellman, 1957) that must be updated with each new sample drawn (Arrow et al., 1949; Furl & Averbeck, 2011; Hauser et al., 2017, 2018; Moutoussis et al., 2011).

Prior work on information sampling has documented that humans sample information sub-optimally, attending to extraneous information (Juni et al., 2016) or through biased weighting of sampling costs (Cisek et al., 2009; Furl & Averbeck, 2011; Hauser et al., 2018). Yet, computing and updating the value of continuing to sample evidence may require significant computational resources, especially for complex decisions, as in sequential information sampling (Bossaerts & Murawski, 2017; Bossaerts et al., 2019; Payzan-LeNestour & Bossaerts, 2011). Indeed, evidence suggests that humans forgo using intensive updating computations, such as Bayesian inference (Charness & Levin, 2005; Gigerenzer & Goldstein, 1996; Payzan-LeNestour & Bossaerts, 2011; Steyvers et al., 2009), even for simpler decisions (Cassey et al., 2016). One factor that might increase the likelihood that humans are willing to expend resources for more optimal computations is the reward value for a correct decision (Bennett et al., 2019; Manohar et al., 2015) but this has not been explored within the context of information sampling. Thus, it remains unclear whether strategies that rely on computations of expected value reflect human information sampling and whether the use of such computations depends on reward context.

If humans do not follow the computations of an optimal decision maker, what determines when they stop sampling? Early accounts proposed that information search relies on simplified heuristic strategies guided by bounded rationality (Conlisk, 1996; Gigerenzer & Goldstein, 1996; Shah & Oppenheimer, 2008; H. A. Simon, 1990; Herbert A. Simon, 1955; Tversky & Edwards, 1966). In these strategies, a set of rules is established to guide both the process of information acquisition (i.e., what information should be attended to and incorporated as evidence) and the decision to stop sampling. Such heuristic strategies minimize cognitive resource expenditure by leveraging declarative rules; however, by definition, these strategies are less flexible and less adaptable to changes in incoming information or changes in context. While recent accounts have demonstrated support for heuristic-style strategies in information gathering (Baumann et al., 2020; Korn & Bach, 2018; Sang et al., 2020), these studies examined information sampling in contexts where individuals received ongoing feedback about their choices, and it remains unclear whether behavior follows heuristic strategies when individuals must collect and integrate information without ongoing feedback – as in the case for many real-world decisions.

In the present study, we investigated whether humans rely on optimal or heuristic strategies (or their combination) during information sampling, and whether their strategies changed as a function of the reward at stake. We tested several potential models of strategic information sampling that varied in the information used and how that information contributed to the decision process. We found that participants’ behavior was best explained by a simple yet flexible strategy in which humans tracked a linear combination of both the evidence in favor of each category and the accrued costs from sampling – but did not rely on a declarative rule or estimations of expected values. This strategy explained a key pattern we observed in sampling behavior: evidence and costs traded off within but not across trials such that as costs accumulated over a trial, the strength of evidence needed for stopping decreased linearly. Moreover, we found that high-reward contexts neither improved optimality nor impacted which strategy best accounted for participants’ decisions. Our results demonstrated how humans implement simple yet flexible information sampling strategies to balance competing motivations for accuracy and cost minimization.

## Results

We tested participants on a modified version of the Information Sampling Task (Fig. 1) (Clark et al., 2006). Participants viewed a series of images randomly drawn from a pool of 25 images. The pool contained images from two categories (e.g., indoor vs. outdoor scenes), with one category comprising 60% of the images and the other comprising 40%. Importantly, participants were unaware of the true proportions of each image category for each trial, although they were told that there would always be a majority category. Participants attempted to identify which category was more prevalent on each trial, under either high ($5.00) or low ($1.00) reward stakes for correct answers. Each image participants chose to draw came with a sampling cost of 2% of the maximum reward on that trial (i.e. $0.10 for a $5.00 trial; $0.02 for a $1.00 trial). Thus, participants had to balance competing goals: sampling more images could increase the accuracy of their guesses, but they would win less reward overall due to the increasing cost accrued (see Supplementary Methods: Task Instructions).

**Figure 1:**
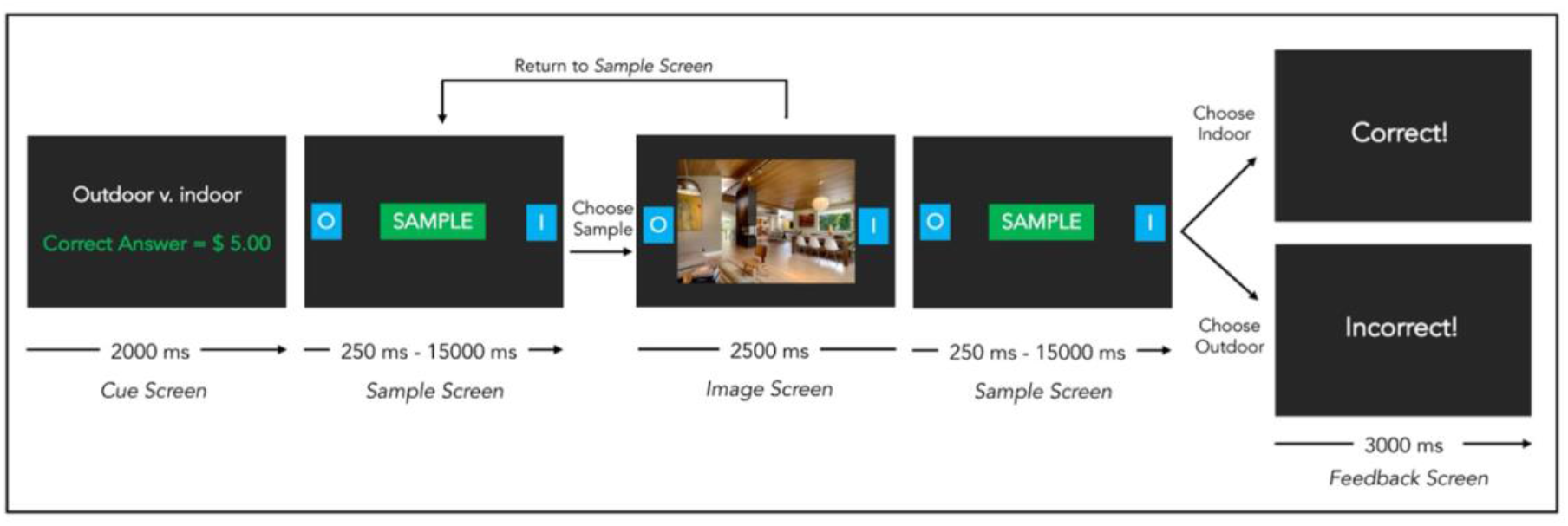
Task Design. Participants completed a modified version of the Information Sampling Task (Clark et al., 2006). Each trial started with a Cue screen that informed the participant which of two domains the images were being drawn from (indoor vs. outdoor or living vs. non-living) and how much money that trial was worth if they performed successfully (high reward: $5.00, low reward: $1.00). Participants were then advanced to the Sample screen where they had a maximum of 15 seconds to sample an image or to indicate their final response for that trial (e.g., O for majority outdoor, I for majority indoor). If they chose to sample, an image appeared over the sample button for 2500 ms. The image then disappeared, and the participant returned to the Sample screen. Each time the participant returned to the Sample screen, the 15 second timer reset. When participants selected a final response, they received feedback on whether they were correct before moving on to the next trial.

### Greater sampling is associated with higher task accuracy, but at the expense of greater cost accumulation

We first investigated how well participants performed the task. Participants correctly identified the majority category 79% of the time (SD = 10%) (Fig 2a). Across the entire task, participants accumulated an average of $95.07 (SD = $10.52) but were only paid for a randomly selected subset of those trials (see Supplementary note 1 and Supplementary Fig. 1). On average, participants viewed 7.82 images (SD = 2.89 images, Range = 1 – 24 images) and reached an average difference in evidence between the currently held majority and minority category of 2.61 images (SD = 0.62, Range = 0 – 8 images) before selecting a majority category (Fig. 2a-c).

**Figure 2.**
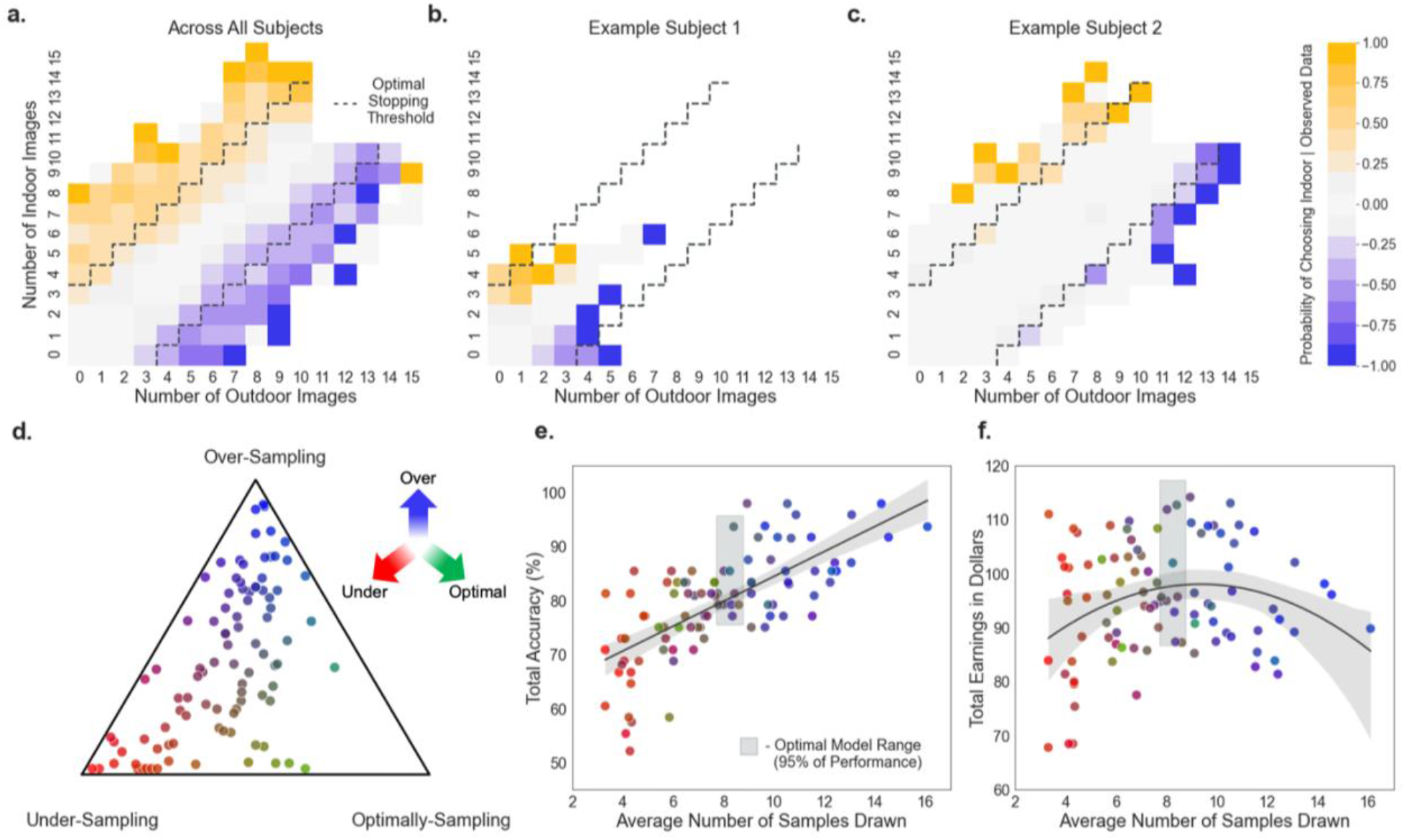
Sampling tendencies between participants relate to overall task accuracy and total earnings. a) The proportion of trials where participants made a stop choice, conditioned on the current combination of images from both categories (collapsed across all participants). For ease of comparison to the task example shown in Figure 1, we label the axes according to “outdoor” and “indoor” categories but note that these data also include behavior from the living/non-living trials as well. The proportion was calculated by dividing the number of instances in which participants stopped by the total number of times all participants reached that combination of evidence. More saturated blues and yellows indicate a higher proportion of stopping across all participants. Dashed grey lines indicate the optimal stopping boundaries. b-c) Probability of stopping given observed data for two example participants. d) Comparison of each participant to optimal behavior. Each trial for each participant was binned into either under-, over- or optimally sampled. The proportion of trials in each bin is represented by the distance to each corner of the simplex and by color. Color mappings for e-f were drawn from d. e) Across participants, the higher the average number of samples a participant tended to draw before stopping, the higher the task accuracy. f) Across participants, the average number of samples predicted total earnings with an inverted-U shaped function. The gray boxes on e and f reflect confidence intervals around the performance range of an optimal model. For plots d-f, each dot reflects a single participant on each graph.

We next examined whether performance outcomes were predicted by participants’ relative tendency to sample. Across participants, a higher average number of samples predicted better task accuracy (Fig. 2e) (F(2, 91) = 43.5, p < 0.001, R^2^ = 0.48, 95% CI [1.47, 3.09]). Similarly, the average number of samples also predicted total earnings via an inverted-U shaped function (Fig. 2f) (F(3, 90) = 3.239, p = 0.026, R^2^ = 0.07, 95% CI (quadratic) [−49.91, −8.39]; *χ*^2^(91, N = 94) = 7.785, p = 0.006). Thus, participants whose sampling was, on average, much lower or much higher than average tended to have lower overall earnings. We repeated this analysis at the trial-level within participants and found consistent results. The number of samples drawn on a given trial predicted both accuracy (*β_samples_* = 0.0818, t = 7.49, p < 0.001, 95% CI [0.06, 0.10]) and earnings (*β_samples_* = −6.49, t = −4.38, p < 0.001, 95% CI [−9.22, – 3.73]; *β*_*sampies*^2^_ = −4.44, t = −3.46, p < 0.001, 95% CI [−6.98, −1.93]). Overall, a higher average number of samples collected prior to stopping resulted in better task accuracy but also greater accumulation of sampling costs – leading to lower earnings.

To compare participant performance to an optimal decision maker (Fig. 2e-f), we computed optimal choices using an Ideal Observer model (see Supplementary Methods: Ideal Observer Model). We used this model to label each trial for each participant as optimal (matched the choice made by the Ideal Observer), under-sampled (stopped sampling earlier than the Ideal Observer), or over-sampled (stopped sampling after the Ideal Observer). We then created composite scores for each participant to assess how close participants were to optimal behavior (Fig. 2d). Collectively, participants performed worse than the Ideal Observer, both in accuracy (two-sided t-test: t(186) = 6.33, p < 0.001, M_ideal_ = 85.6%, SD = 5.2%) and in earnings (two-sided t-test: t(186) = 6.02, p < 0.001, M_ideal_ = $101.97, SD = $7.67), but they did not differ significantly from the Ideal Observer in the average number of samples drawn (two-sided t-test: t(186) = 1.38, p = 0.169, M_ideal_ = 8.24 images, SD = 5.21). Upon further examination, however, this was due to participants either over- or under-sampling relative to optimality (F(5, 530) = 20.54, p < 0.001), indicating that participants are either estimating optimal behavior poorly or relying on a different process to establish stopping criteria.

### Models of information sampling strategies

Our next analysis investigated how participants sampled and integrated information towards a decision. We first identified four potential information sampling strategies that relied on a range of optimal and heuristic approaches (see Supplemental Methods: All Sampling Strategies). Our first strategy was a probabilistic modification of the optimal Ideal Observer (see Supplementary Methods: Expected Value Model Formulation). This strategy relied on the same Bayesian updating and inference to estimate the expected value of each option but was adapted to allow for inherent noise in participant decision making as well as to account for cost accrual mechanisms that deviated from objective costs accumulation (Cisek et al., 2009; Ditterich, 2006; Furl & Averbeck, 2011; Hauser et al., 2018) (see Methods). The cost mechanism that best accounted for participant behavior was similar to the strategy from Hauser et al., (2018), in which subjective costs were accumulated nonlinearly, representing a growing urgency across information sampling to select a final option (Cisek et al., 2009). As such, the Expected Value Urgency Threshold (EV-UT) strategy predicted that participants would sample until the value of an option surpassed the value of continuing to gather information, and that the value of continuing would sharply decline after the urgency threshold was met (Fig. 3, EV-UT).

**Figure 3.**
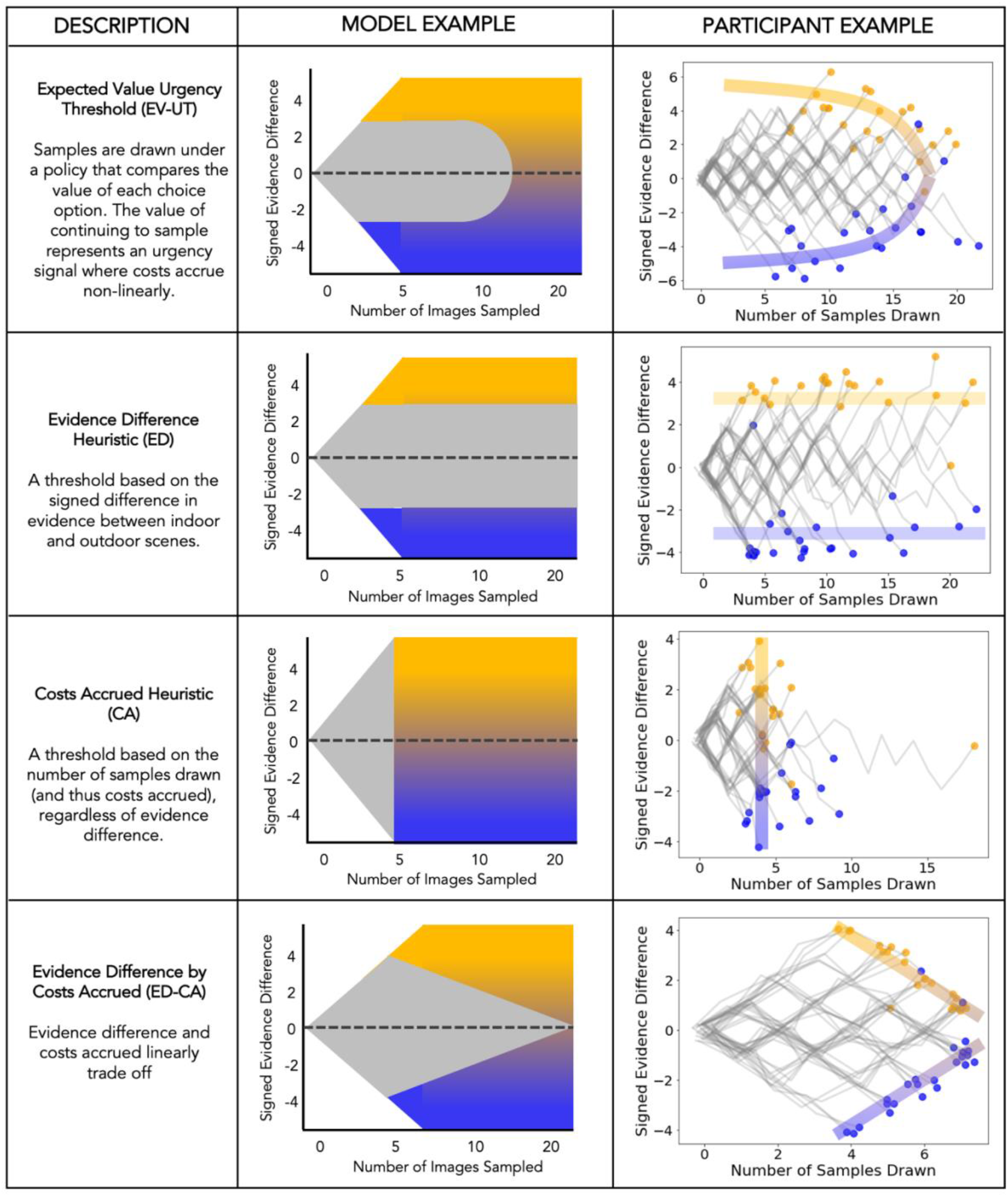
Tested information sampling strategies with model predictions and example representations. We hypothesized that information sampling patterns would represent one of four potential strategies. The left column indicates the description of the sampling behavior for each hypothesized strategy. The middle column reflects an example model prediction for choices given an example set of parameters. Gray portions indicated predicted choices to continue collecting information where yellow and blue areas indicate predicted choices for selecting a particular majority category (e.g., indoor or outdoor). More saturated areas of yellow and blue reflect choices with strong evidence for that choice while less saturated areas reflect choices for which the evidence is weaker. The last column represents examples of each strategy seen in our participant data. Gray lines indicated participant choices to continue sampling and dots represent the decision to select a majority (yellow = choose indoor, blue = choose outdoor). Colored bars are illustrative to exemplify each strategy.

The next two strategies were probabilistic adaptations of two common heuristics. The first, Evidence Difference Heuristic (ED), was a heuristic that assumed participants tracked the continuous signed difference in evidence between the two categories towards a threshold (e.g., “I sample until one category has 4 more than the other”). This strategy predicted that participants approach information sampling insensitive to the number of images sampled and implies that the stopping boundary is stationary and constant across sampling (Fig. 3, ED). The second, Costs Accrued Heuristic (CA), was a heuristic that assumed, participants used the continuous number of samples drawn as a proxy for the costs accrued from sampling and only a binary representation of the difference in evidence to inform choice (e.g. “I sample 5 images and then choose the majority). Similar to the first, this strategy predicted a stationary threshold that triggered the end of sampling, but now, bound to the number of samples drawn, implying that the magnitude of evidence mattered less (Fig. 3, CA).

The last tested strategy, Evidence-by-Costs Tradeoff (ED-CA), was a combination of the two heuristic approaches, such that participants used continuous representations of both the difference in evidence between the categories and the number of images collected to inform their choices. These two quantities were then linearly combined towards a threshold. This strategy predicted that as the number of samples collected increased, the difference needed between the evidence in favor of one category over the other would decrease, representing a non-stationary but constant tradeoff between the two informational sources (Fig. 3, ED-CA).

Across these four strategies, there were striking visual distinctions in the types of choices each predicted (Fig. 3, Model Example). Furthermore, models were distinguishable from one another, as shown by model recovery such that choices generated by each model were best fit by the model that generated them (see Supplemental Methods: Parameter Recovery and Model Recovery). Thereby, we were not only able to detect descriptive differences in sampling strategies but were also able to draw inferences about the underlying process guiding information sampling (Fig. 3).

### Moment-by-moment sampling decisions were best predicted by an Evidence-by-Costs (ED-CA) strategy

We next fit each participant’s data to each of the four sampling strategies outlined above. Participants overall were best fit by the Evidence-by-Costs Tradeoff (ED-CA), which used continuous representations of both the difference in evidence between the categories and the number of images collected to inform their choices. This strategy outperformed the three other proposed strategies such that of our 94 participants, 83 were best fit by the ED-CA strategy (Fig. 4a). A repeated-measures ANOVA confirmed that the ED-CA strategy had significantly lower BIC scores (F(3, 279) = 81.17, p < 0.001) compared to the CA strategy (t(93) = −14.65, p < 0.001), the ED strategy (t(93) = −10.92, p < 0.001), and the EV-UT strategy (t(93) = −11.32, p < 0.001) (Fig. 3b-d). There was no difference in fit between the ED strategy and the EV-UT strategy (t(93) = 1.17, p = 0.24). The CA strategy fit the worst of all four strategies tested (ED and CA: t(93) = −4.83, p < 0.001; EV-UT and CA: t(93) = −5.40, p < 0.001).

**Figure 4.**
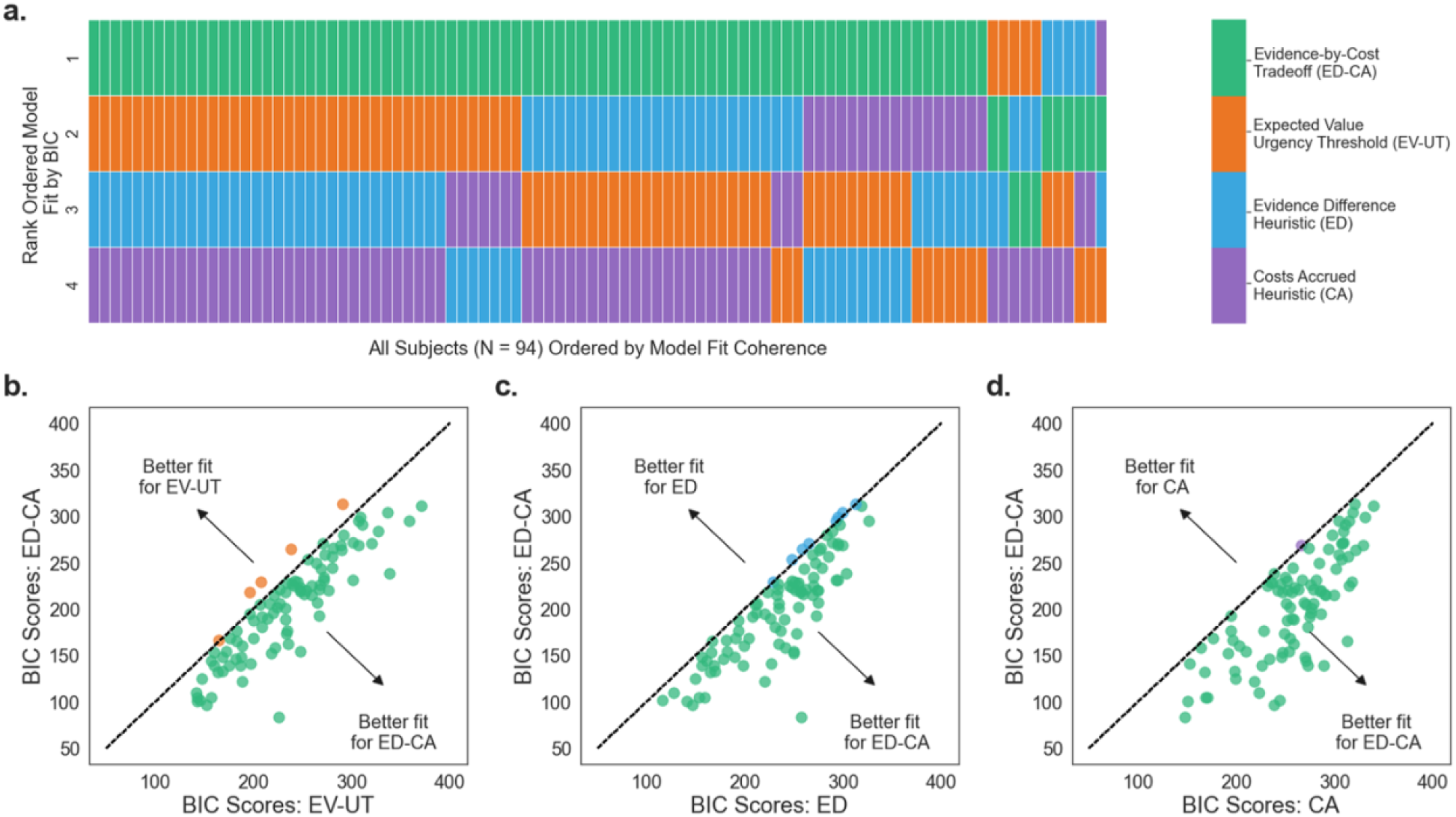
Evidence-by-Costs (ED-CA) strategy outperforms both simpler heuristics and expected value models in predicting decisions. BIC scores were calculated for each participant under each model. a) We ranked BIC scores from lowest (better fit) to highest (worse fit). Each participant represents a column on the heatmap. A rank of 1 indicates the best fitting model for that participant. Green represents the Evidence-by-Costs (ED-CA) strategy, orange represents the Expected Value Urgency Threshold (EV-UT) strategy, blue represents the Evidence Difference Heuristic (ED) strategy, and purple represents Costs Accrued Heuristic (CA) strategy. b-d) Plotted BIC scores for the ED-CA strategy compared to all other strategies. Each dot reflects a single participant. Dots are color coded by which model had the lower BIC score between the two models. b) BIC scores plotted between ED-CA and EV-UT. c) BIC scores plotted between ED-CA and ED. d) BIC scores plotted between ED-CA and CA.

Results from the ED-CA strategy revealed that participants were sensitive to the costs of increased sampling (i.e., number of samples drawn) and the magnitude of evidence favoring one category over the over (i.e., signed difference in samples). We next sought to test whether there was a linear tradeoff between the difference in evidence for each category and the number of samples.

### Stopping criteria were non-stationary within a trial but stationary across trials

The Evidence-by-Costs (ED-CA) strategy aggregates the currently available information and compares it to a decision threshold that reflects a participant-specific level of information needed to terminate sampling. A prediction of this strategy is that, over the course of a trial, the contributions of total samples drawn and evidence difference can trade off against each other. We found that the number of samples drawn negatively predicted evidence difference at the time of stopping (*β_samples_*: −0.0722, t(66.6) = −6.93, p < 0.001 CI [−0.09, −0.05]) and that the majority of our participants (N=78/94) evinced this negative relationship (Fig. 5). This result indicates that the more images sampled (and thus the higher the costs), the less the evidence difference must be in order to stop sampling, consistent with accounts of collapsing boundaries in evidence accumulation (Ditterich, 2006; Drugowitsch et al., 2012; Malhotra et al., 2017; Murphy et al., 2016). We tested whether this relationship was better fit with the inclusion of a quadratic term, but that inclusion did not significantly improve model fit (*χ*^2^(1, 94) = 2.94, p = 0.09). Interestingly, the initial threshold for stopping is similar to that of the Ideal Observer model but quickly decreases as the trial progresses.

**Figure 5.**
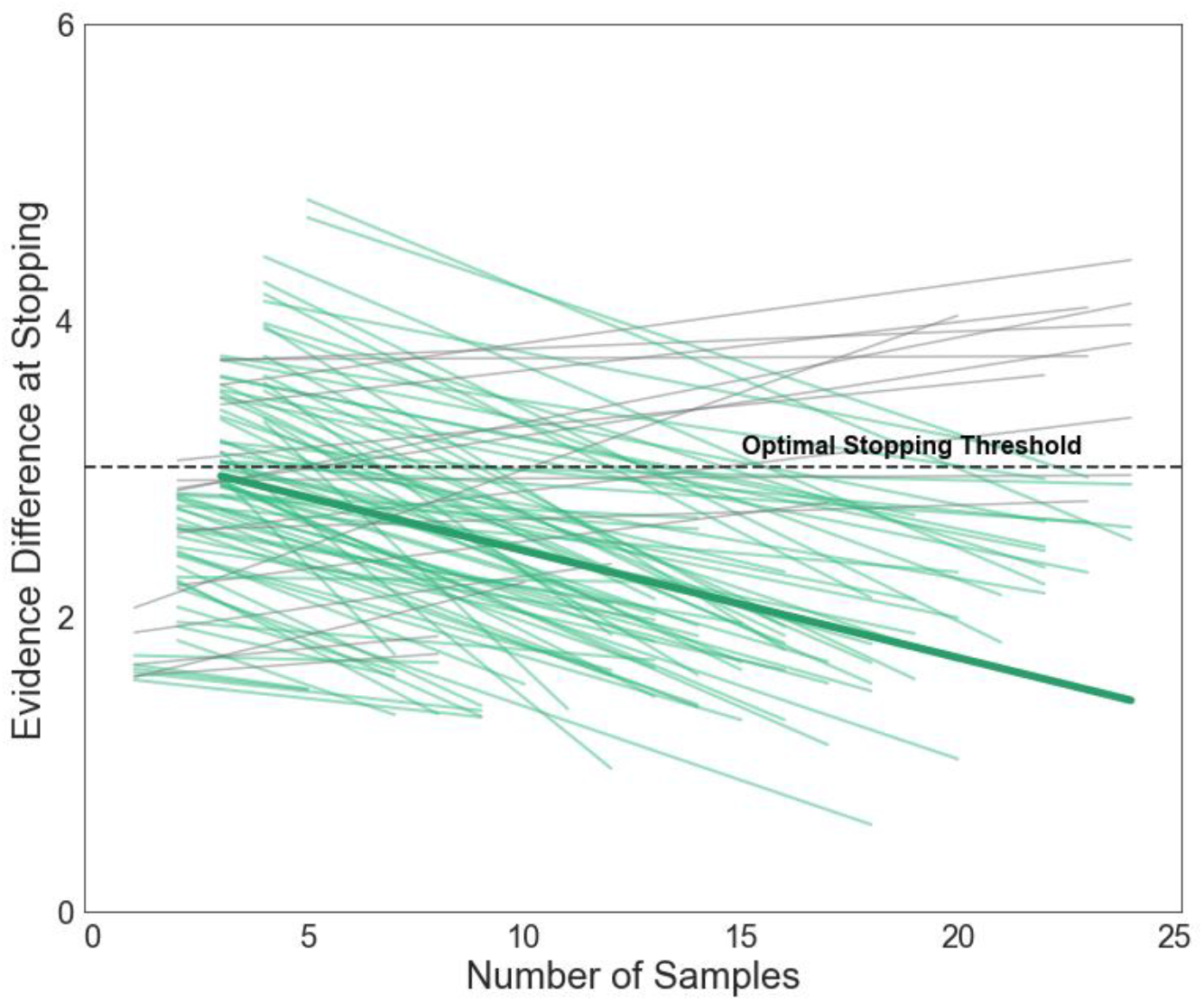
Evidence difference and number of samples trade off. Individual slopes modeled through linear mixed effects model. As the number of samples increased, the difference in evidence needed to stop sampling decreased. Green lines represent participants with negative slopes (N=78) and grey lines represent participants with slopes greater than or equal to zero (N=16). The mean slope is represented by the bold green line. The optimal stopping threshold from the Ideal Observer, which stays constant throughout sampling, is represented as the dotted black line. Lines are truncated to reflect the minimal and maximal number of samples prior to stopping for a given subject. Estimates for each participant and the mean are taken from linear mixed effects model with maximum random intercept and slopes.

We also examined whether stopping criteria changed across trials in the task. We first investigated whether there were changes in stopping criteria (i.e., changes in participant’s average sampling number or evidence difference prior to stopping) or changes in stopping consistency between the first and second half of the task (i.e., changes in the standard deviation of samples drawn or evidence difference). None of the dependent measures were significantly different from the first to the second half of the experiment (all p > 0.05). Additionally, there were no significant differences in accuracy from the first to the second half of the task (p > 0.05), indicating that performance was relatively stable from trial to trial. Moreover, none of the dependent measures were significantly different as a function of the outcome of the previous trial (all p > 0.05). Collectively, these results suggest that behavior in this task was stable across trials.

### Increasing reward stakes does not encourage use of more optimal strategy

Lastly, we investigated whether an increase in reward stakes shifted strategy use across our participants. We hypothesized that under high reward contexts, participants might expend more computational resources to sample more optimally (Bennett et al., 2019; Manohar et al., 2015), thus participants’ behavior would be more similar to the Ideal Observer or be better fit by a strategy that relies on optimal value estimation (e.g., the EV-UT strategy). To test this, we split trials by reward condition within each participant and fit each subset of behavior to the ED-CA strategy and to the EV-UT strategy. We found that overall, the ED-CA strategy provided the best fit for the majority of our participants (N=69) across both high-reward and low-reward contexts. Of the participants that had different fits between high-stakes and low-stakes trials, there was not a consistent pattern of change (EV-UT strategy provided a better fit under high-reward trials for 7 participants and provided a better fit under low-reward trials for 12 participants) (See Supplementary Note 4, Fig. 3 for more detail).

Similarly, changes in sampling as a function of reward condition did not encourage optimal behavior compared to the Ideal Observer. We found no difference in the proportion of trials in which participants sampled optimally across reward conditions (t(441) = 0.847, p = 0.958). Instead, we see an increase in the relative number of over-sampling trials in high-reward contexts, compared to low-reward contexts (t(446) = 3.068, p = 0.023) (Fig. 6a). Follow-up trial-by-trial analyses confirmed that participants sampled more images under high-reward compared to low-reward (*β_reward_* = 0.503, t = 4.762, p = < 0.001 CI [0.29, 0.71]) (Fig. 6b) and required a larger evidence difference to stop (*β_reward_* = 0.147, t = 4.428, p < 0.001 CI [0.08, 0.21]) (Fig. 6c). Despite the adjustments in sampling behavior under high-reward compared to low-reward, there was no difference in performance outcomes between the two conditions (*β_reward_* = 0.02, t = 0.25, p = 0.803). High-reward trials, thereby, did not encourage more optimal choices but instead, drove participants to simply adjust the threshold for stopping within their existing strategy. These adjustments, however, ultimately did not improve performance on high-reward trials. Our reward manipulation provides support that participants adopt a global policy for stopping behavior – and adjust that policy in different reward conditions – but do not systematically deploy different policies in different reward contexts.

**Figure 6.**
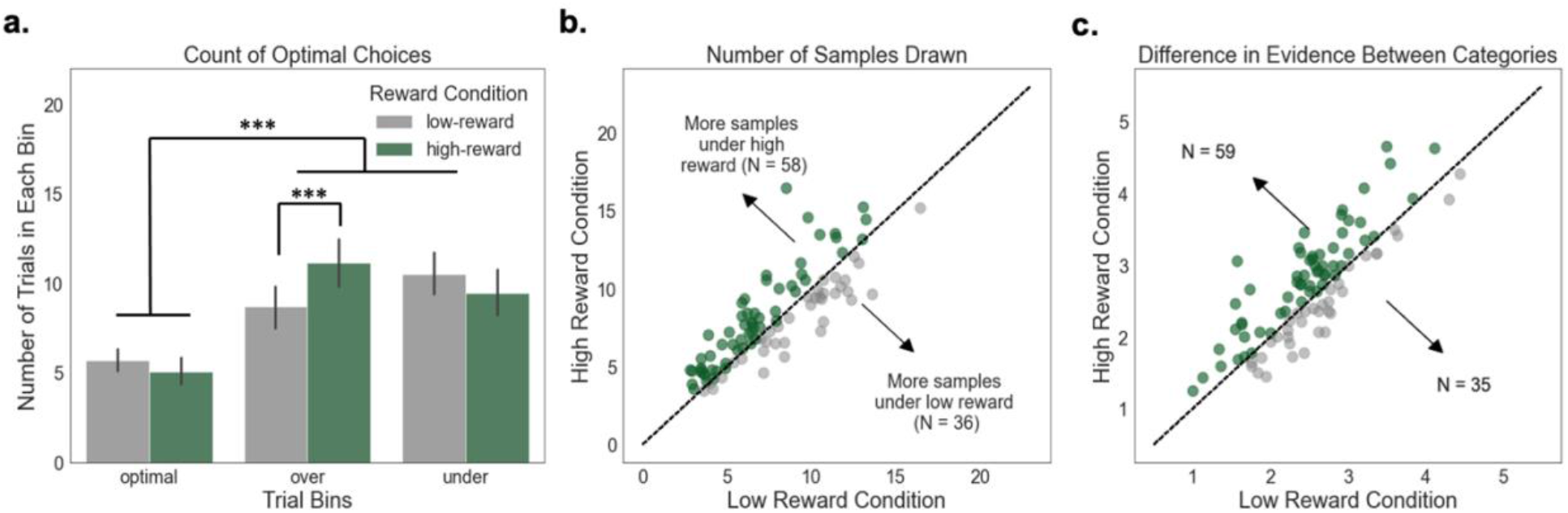
High reward stakes increase sampling but do not encourage more optimal behavior. Changes in behavior as a function of reward condition were assessed across three variables of interest: changes in the proportion of optimal choices, number of samples drawn, and the evidence difference at time of stopping. a) Both over- and under-sampling were significantly more common than optimal sampling. We also observed an interaction such that in high-stakes trials, participants over sampled more than in low-stakes trials. Error bars reflect standard error of the mean. b) Participants sampled more images under high stakes compared to low stakes. c) Participants waited for greater evidence differences in high-stakes trials compared to low-stakes trials. Each dot represents a participant; color coding indicates the condition with greater sampling. Dots closer to the diagonal indicate less change between conditions, while dots that deviate from the line indicate greater changes by reward condition.

## Discussion

Balancing the costs of gathering information with the desire to accurately choose outcomes represents a fundamental challenge for human decision-making. Various theoretical models of information sampling have been developed to explain how humans address this challenge, but these models tend to either emphasize resource-intensive optimal computations or efficient but rigid heuristics. Our results support an alternative perspective: information sampling relies on a computationally simple and flexible strategy that accumulates task-relevant information until a threshold is reached. This Evidence-by-Costs strategy also predicted other key features of sampling behavior, including the tradeoff such that more evidence was required when costs were low but that less evidence was required when accrued costs were high. In addition, we found that changing the reward value for a correct choice did not encourage more optimal behavior nor a switch to using more computationally expensive estimations. Collectively, we concluded that humans flexibly adapt the information accumulation process to balance competing motivations for accuracy and cost minimization.

Previous work has demonstrated that humans exhibit subjective biases in the computation and integration of accruing costs that limit the optimality of behavior (Cisek et al., 2009; Coenen & Gureckis, 2016; Furl & Averbeck, 2011; Hauser et al., 2017, 2018). However, these policies still assume that humans engage in the computational expensive estimations of the value of stopping and the value of continuing in the same manner as an Ideal Observer (Bossaerts & Murawski, 2017; Payzan-LeNestour & Bossaerts, 2011). The Evidence-by-Costs strategy provides a starkly simpler rule that reduces computational demands through limits both on the number of steps involved and the amount of information held in memory (Bossaerts & Murawski, 2017; H. A. Simon, 1990). It also provides a better account of behavior: instead of computing the probability of success in accordance with Bayesian inference, participants relied on a mechanism that estimated success by tracking the evidence difference between the two categories of images.

The use of an Evidence-by-Costs strategy was evident in the trial-by-trial behavior of our participants. Consistent with prior work (Ditterich, 2006; Malhotra et al., 2017; Murphy et al., 2016), we found that there was a tradeoff between two quantities: as the number of samples (costs accrued) increased, the smaller the evidence difference needed to stop sampling. The existence of this tradeoff could reflect two possibilities. First, as more samples are drawn, individuals could be relying on smaller but more reliable differences between the majority and minority category. Although we cannot rule this possibility out completely, it is unlikely given that participants were kept blind to the true underlying distribution and thus the proportion of the samples not yet drawn was also unknown. Alternatively, this tradeoff could mean that the stopping threshold may be reached from different linear combinations of two quantities: a large number of samples or a large evidence difference, and individuals actively use both to inform stopping. The result is that as sampling continues and costs are accrued, people begin to prioritize different information – moving from an initial bias toward larger evidence differences toward a later bias against costs.

Our results extend prior research positing tradeoffs between evidence and sampling costs, akin to the speed-accuracy tradeoff, to determine choice. Murphy et al. (2016) saw this tradeoff under conditions in which participants were pressed for time, yet our results suggest that this trade-off can exist even without external time pressures. Additionally, Malhotra et al. (2017) found that the tradeoff between costs and evidence only existed when participants completed intermixed trials of varying difficulty, yet we show that the trade-off persists even when difficulty is fixed across trials. It is possible that our task, similar to that of Malhorta et al. (2017), had increased uncertainty because the distribution of images from either category was unknown to our participants. This could have encouraged a strategy that did not rely on a single information factor (e.g., evidence) to inform stopping. Additionally, our tradeoff appears linear in form, as opposed to exponential or sigmoidal, which are indicative of a growing urgency signal (Cisek et al., 2009). Unique to our results is that this tradeoff was predicted by the specific strategy (Evidence-by-Costs) individuals used to arbitrate between sampling and stopping.

We also found that increasing the financial stakes for a trial did not encourage adoption of a strategy that relies on optimal value estimation nor did it increased the likelihood of optimal behavior (i.e., participants did not move closer to the stopping decisions that would be made by an Ideal Observer (Achtziger et al., 2015)). Instead, we saw that participants were still best fit by the Evidence-by-Costs strategy. Behaviorally, participants in the high-reward condition over-sampled, effectively raising the stopping threshold without changing the underlying computations; both drawing more samples and waiting until they achieved a larger difference between categories. This stands in contrast with prior work that suggested increasing monetary stakes can increase one’s motivational state, thereby encouraging the use of more optimal but expensive strategies (Bennett et al., 2019; Manohar et al., 2015).

Why might increased monetary stakes encourage behavioral adjustments but not push participants towards optimal behavior? One possible explanation is that conditions of equal difficulty but with higher monetary stakes could increase the effort put into the trial because of lower opportunity costs (Otto & Daw, 2019; Shenhav et al., 2013), but still not warrant the adoption of a completely new and expensive strategy. This is likely the case given that our task had intermixed high and low reward trials, such that participants would have to continually switch between strategies, which introduces cognitive costs (Luwel et al., 2009). Alternatively, our task was more difficult than prior information sampling paradigms because participants had to remember their previous samples – and thus the added memory demands might have deterred adoption of a more resource-intensive strategy. Although prior work has suggested that working memory capacity can impact the amount and use of information during information acquisition (Rakow et al., 2010), we did not measure the working memory capacities of our participants. Future research will need to explore how working memory demands and the cost of switching information sampling strategies shape stopping policies.

This study raises an important set of questions regarding how individuals determine their idiosyncratic thresholds. Evidence-by-Costs sampling provides two unique informational components that contribute to stopping – and individual model fits revealed variability in how participants weighed information about evidence and costs. Specifically, variability in the weighting of evidence could reflect varying levels of confidence required for stopping across individuals, as seen in other work (Hausmann-Thürig & Läge, 2008). In line with prior work (Hauser et al., 2017, 2018; Juni et al., 2016; Otto & Daw, 2019; Petitet et al., 2021), we also see individual differences in sensitivity to accruing costs. Future work towards encouraging more optimal behavior can leverage our approach by specifically targeting informational components that most contribute to a person’s sub-optimal sampling behavior. For example, an individual who consistently over-samples might do so because they are more sensitive to accuracy (a higher threshold starting point) or because they are less sensitive to accruing costs (a shallower trade-off slope). The ability to arbitrate between potential sources of error could provide a more targeted prescription to ameliorating the cause of over-sampling.

Moreover, our findings emphasize key directions for understanding sampling strategies themselves. First, additional research should identify and delineate strategies that do not completely conform to either heuristics or optimal behavior. A recent study (Korn & Bach, 2018), demonstrated the use of both heuristic and optimal strategies (but not a combination of the two) across a foraging task, providing insights into factors that shape strategy selection; for example, higher levels of experienced uncertainty may push sampling toward optimality. Similarly, in our current study, the Evidence-by-Costs strategy did not specifically integrate components from the optimal strategy but was sensitive to the same information sources. Cataloging a more complete space of sampling strategies will advance our understanding of how humans select what information to attend to and how that information is transformed into potential actions.

Future research should also explore how people determine what strategies to implement in different contexts. During information sampling, individuals not only decide how to balance sampling costs with accuracy but also contend with balancing the costs and benefits of exerting control (Shenhav et al., 2013). Previous information sampling accounts have examined the impact of contexts such as changes in task difficulty (Coenen & Gureckis, 2016; Malhotra et al., 2017) and changes in sampling costs (Hauser et al., 2018; Juni et al., 2016) in altering sampling behavior but have not specifically examined if these contexts changed the underlying strategy. We examined the context of varying reward stakes on information sampling and found that while individuals maintained the same underlying strategy between both contexts, reward increased the overall information that people gathered. Our results differ from that observed in a reinforcement learning task, where reward stakes resulted in a switch to a more intensive but optimal strategy (Bennett et al., 2019). Additional investigation is needed into how individuals use context to evaluate when to switch information sampling strategies and when to adapt an existing strategy. Prior work has indicated that humans can learn to use context to determine strategy selection (Lieder & Griffiths, 2017; Payne et al., 1988; Rieskamp & Otto, 2006); for example, experience with a problem leads to adoption of more heuristic strategies and can even direct selection amongst different heuristics (Rieskamp & Otto, 2006). Although we did not find any changes in strategy in the current study, our task involved a longer sampling process and fewer sampling episodes, thus making it harder for participants to explore a variety of strategies. Future work will need to investigate how much experience individuals need in order to use contextual factors to inform both the selection and implementation of information sampling strategies.

Information sampling is a complex but ubiquitous challenge for decision makers. In the present study, we show that humans confront this challenge by adopting a strategy that balances the efficiency of heuristics but with increased flexibility. Specifically, our results demonstrate that unlike optimal strategies, humans are performing simple accumulations instead of computing expected values, and unlike heuristic strategies, humans are dynamically integrating information instead of using rigid rules. Future work expanding how humans build such flexible strategies and how individual differences determine the relative weighting of different elements of those strategies (e.g., reward sensitivity) will provide further insight into the mechanisms by which bounded rationality guides decision-making processes.

## Methods and Materials

### Participants

Participants (N = 105, Mean age = 26.14, SD = 4.79, 69 female) were recruited from the Durham community using flyers and online postings. Our demographic breakdown included 37 participants who identified as White/Caucasian, 47 identified as Asian/South Asian, 14 identified as Black/African American, 3 identified as Hispanic/Latinx, and 4 identified as multi-racial/ethnic. To participate, individuals had to 1) be within the age range of 18-50 years old, 2) have no history of neurological injury or disorders (including seizures and epilepsy), and 3) be fluent in English. Eleven participants were excluded from all analyses, three due to computer error and eight due to having unusable sampling data (failed to sample more than once on over 25% of trials), leaving a final total of 94 participants. All participants received informed consent under the guidelines of Duke University’s Institutional Review Board.

### Procedure

At the outset of each experimental session, participants provided informed consent, received task instructions (see Supplementary Methods: Task Instructions) before practicing the experimental task (Fig. 1). Participants returned to the laboratory approximately 24h later for to complete a surprise memory test for the images sampled during the first experimental task. Results for the memory test can be found in the Supplement (see Supplementary Methods: Memory Task, Descriptions, and Findings) but will not discussed in the main manuscript.

Participants performed a modified version of the Information Sampling Task (Clark et al., 2006) displayed using PsychoPy 2.7 (Peirce et al., 2019). Participants were told that on each trial there was a box that contained 25 images from one of two possible domains: scenes and objects. Each domain had two categories (scenes: indoor or outdoor, objects: living or non-living) and each image belonged exclusively to one category. Each trial contained images from only one domain. Images were all naturalistic photos collected from Google Image searches and scaled to the same size in pixels.

On each trial, participants were tasked with identifying the underlying majority category for a given domain. Participants could sample images from the box serially until they felt they had enough evidence to select a majority category (max of 25 images per trial). Participants were told that there would always be a majority category, but they were not told the true proportions of each image category and were instructed that the proportions could change between trials. The true proportion was kept constant at 60/40 for majority/minority categories (i.e., 15 of the 25 images would be from the majority category). The order of the images was randomized. Participants performed trials under high ($5.00) and low ($1.00) reward stakes. Incorrect responses in both stakes conditions resulted in a reward of $0.00 for that trial. In addition, participants incurred a cost for each sample they made (2% of the max reward they could earn for that trial). Thus, participants had to balance their confidence in identifying the true majority against accruing sampling costs.

At the start of each trial, a *cue screen* (2000 ms) appeared, informing participants of the image category judgment (e.g. indoor vs. outdoor or living vs. non-living) as well as the monetary reward available for a correct response (e.g. Correct Response = $1.00/Correct Response = $5.00, before sampling costs). They then viewed the *sample screen,* whereupon they had the option to either sample an image or make a final choice as to what category they thought predominated on the trial. If they chose to sample (by selecting the down arrow key), one image would immediately appear in the middle of the screen for 2500 ms *(image screen).* After the image disappeared, participants were returned to the *sample screen*. Images did not stay visible to participants after the 2500 ms presentation; thus, participants had to remember past images to guide their choices. At each instance of the sample screen, participants had 15 seconds to make a choice before they automatically advanced to the next trial, with the previous trial being marked as incorrect. This happened on approximately 0.003% of trials across all participants (17 out of 5004 trials).

Participants were free to sample as few or as many images as they deemed necessary to guess the more prevalent category. When participants decided to stop sampling, they indicated their decision about which category they felt predominated on that trial by choosing the box (by pressing either the right or left arrow key) that was associated with that category, which were displayed on either side of the sample button throughout the trial. After participants made their final choice, they were shown a *feedback screen* (2000 ms) that displayed if their guess matched the true majority in the box (e.g. “Correct”/ “Incorrect”).

Participants completed 48 trials in the task. Trials were fully counterbalanced such that they saw an equal number of trials from either category, and each category was equally represented in both high and low reward stakes. Additionally, each category had the same overall probability of winning. To ensure incentive compatibility, participants were paid for 4 trials, randomly chosen. Because the task was self-paced and participants varied in how many images they collected, the session length ranged from 11 minutes to 47 minutes (Mean time: 24.86 minutes, SD: 7.63 minutes).

### Data Analysis

To understand how participants determined when to switch from gathering information to selecting a final choice, we compared participants’ behavior using a series of computational models. We first measured how close each participant’s stopping choices were to the Ideal Observer (model-predicted optimal choices). We then fit each subject’s behavior to four sampling strategies. The first strategy, Expected Value Urgency Threshold (EV-UT), relied on expected value computations to inform choices. We used an adaptation of this strategy similar to Hauser et al., (2018), that suggested humans integrate costs non-linearly. In this strategy, the threshold to transition from sampling to selecting an option was both non-stationary and inconstant across the number of images collected. The second strategy, Evidence Difference Heuristic (ED), was a heuristic that assumed participants tracked the continuous signed difference in evidence between the two categories towards a threshold (e.g., “I sample until one category has 4 more than the other”). This strategy suggests that participants approach information sampling insensitive to the number of images sampled and implies that the stopping boundary is stationary and constant across sampling. The third strategy, Costs Accrued Heuristic (CA), was another heuristic that assumed, participants used the continuous number of samples drawn and only a binary representation of the difference in evidence to inform choice (e.g. “I sample 5 images and then choose the majority). Similar to the first, this strategy maintained a stationary threshold that triggered a decision to select an option but implied that the magnitude of evidence mattered less. The last strategy, Evidence-by-Costs Tradeoff (ED-CA), was a combination of the two heuristic approaches, such that participants used both continuous representations of the difference in evidence between the categories and the number of images collected to inform their choices. This strategy reflected a linear threshold that decreased as the number of samples collected increased, representing a non-stationary but constant tradeoff between the two informational sources. Detailed descriptions of the strategies are outlined below.

In all models, choices were assumed to be probabilistic and were all fit using a SoftMax function. To emphasize, participants were given the following information: each box on each trial contains a total of 25 unique images, the maximum reward value for a trial is either $5.00 or $1.00 and the cost per image is a constant 2% of the maximum reward available on a trial ($0.10 for $5.00, $0.02 for $1.00), the proportion of images from either category is specifically withheld and participants are told that the proportion may change on a trial-by-trial basis.

#### Optimality

To compare participant sampling behavior to that of an Ideal Observer, we first calculated the optimal stopping points using a model adapted from Hauser et al. (2018). Because the proportion of reward to costs was equivalent for high vs. low stake trials, the computations and optimal stopping points are the same across reward conditions. After each sample (*N_samp_*), the optimal agent compares the value of stopping given the current evidence against the value of continuing to sample. In order to determine the value of each action, the agent computes the probability of success in selecting the correct category given the current evidence (i.e., the number of indoor (*n_i_*) and outdoor samples (*N*|*samp* — *n_i_*) collected thus far). Because the true underlying distribution of indoor to outdoor images is unknown, the optimal agent must also estimate the underlying distribution *(q)* from which the samples are being drawn from. Then, it must compute the probability of success under each possible proportion of majority to minority images weighted by the likelihood that that is the true distribution (Eq. 1.1, 1.2). We set prior beliefs, *α* and *β*, about the true underlying distribution equal to 1.

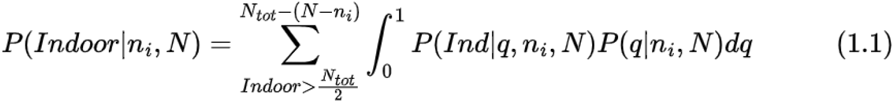

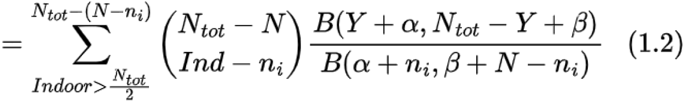

The expected value of stopping is then computed by taking the probability of success of stopping multiplied by the reward ($5.00 for high stakes, $1.00 for low stakes) minus the accrued costs, *c*, per sample ($0.10 for high stakes, $0.02 for low stakes) (Eq. 2).

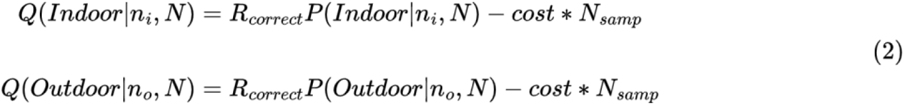

The expected value of stopping is then compared to the expected value of continuing to sample. To compute the expected value of continuing to sample, the optimal agent calculates the expected value of stopping for each state using backward induction to solve for the Bellman equation (Bellman, 1957). Briefly, the expected value of continuing at timepoint 25 is equal to 0 because no additional samples can be drawn. Thus, the expected value at timepoint 25 is equal to the expected value of stopping given all available evidence. Given a behavioral policy that always chooses the highest valued action, the value of all possible states at timepoint 24 (and prior timepoints) can then be calculated using backward induction. Thus, for each possible state, the expected value of continuing, averages over all potential future states, weighting them by the likelihood that that state will be reached (Eq. 3). *s*’ represents the next immediate state, which can either reveal another indoor image (*i* = 1) or an outdoor image (*i* = 0).

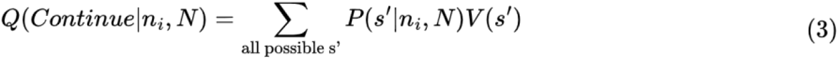

To examine how participants’ behavior compared to optimal behavior, we binned each trial for each participant as either optimal, under-sampled, or over-sampled based on where each stopping decision fell compared to optimal. Because of the cost and reward structure of this specific task, optimal behavior followed an easily verbalized heuristic of “sample until a difference of 3 is achieved.” This heuristic fits with “fast and frugal” criteria of being computationally simple and relying on only a fraction of available information but still preforming optimally (Gigerenzer & Goldstein, 1996; Todd & Gigerenzer, 2000). Thus, optimal behavior could be achieved through multiple routes of computation.

#### Expected Value Computation Strategy

We examined a probabilistic modification of the optimal strategy. This strategy relied on the same Bayesian updating and inference to estimate the probability of success given the available evidence but was adapted to allow for inherent noise in participant decision making as well as to test different cost accrual mechanisms (see Supplementary Methods: Expected Value Model Formulation). Prior research has documented that human deviation from optimality could arise from the accumulation of costs that are different from the specified objective sampling costs (Cisek et al., 2009; Ditterich, 2006; Hauser et al., 2017, 2018). We therefore tested whether the cost per step (*c_step_*) was being subjectively accrued in either a linear (Eq. 5), or non-linear (sigmoidal, Eq. 6) manner, and if these outperformed the use of objective costs (Eq. 4). In equations 5, *t* represents the subjective scaling of objective costs. In equation 6, p represents the sample number where costs begin to accumulate. In all equations, *R* represents the reward condition, 0.02 represents the percentage of the max reward, which equates to the objective cost per sample. Overall, our non-linear cost accrual outperformed our other two models of cost (see Supplementary Methods: All Sampling Strategies).

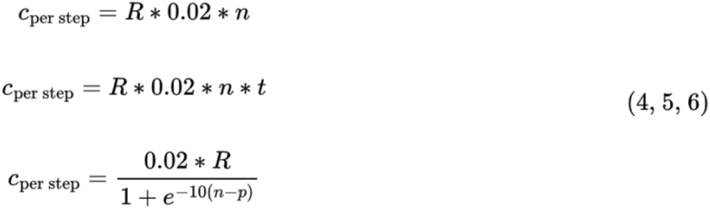

To test the different models of cost, we isolated the impact of costs to the choice to continue sampling. To do so, we updated the action values for choosing each final option as well as the value of continuing to sample as such.

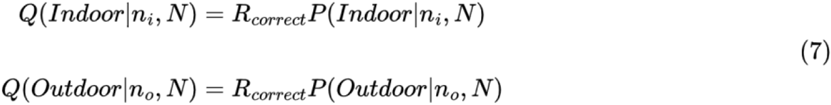

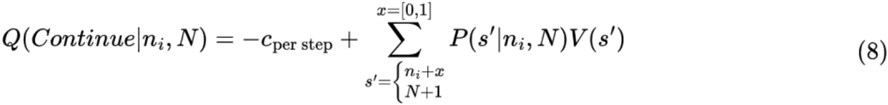

These expected values were then transformed into probabilities using the following Softmax function with inverse temperature parameter, *β*, and irreducible noise parameter, *ξ* (Eq. 9). Importantly, this first family of models relied on the assumption that humans were still performing the underlying Bayesian operations to determine their policies, albeit with noise in their choice process.

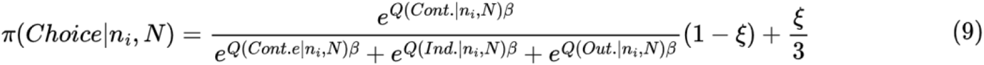

For all of the models tested within the Expected Value Computation framework, participant data was best fit by the subjective non-linear cost model, giving rise to the Expected Value Urgency Threshold strategy and replicating previous work (Hauser et al. 2018). Given our two reward contexts, we also tested whether participants adapted this strategy based on the reward available for that trial. To do so, we tested three separate modifications of the subjective non-linear Expected Value Computation strategy. In our first model, we fit separate models for each reward condition for each participant, suggesting that participants could have completely difference parameter values for each reward condition. In our second model, we fit one model for both reward conditions and included a parameter that scaled the reward value for low-reward trials to be between $1.00 and $5.00, suggesting that the parameter values for both conditions could be equivalent, but participants were still sensitive to the difference in reward outcomes. Our last model either through the same model under just one of the t the high reward condition models treated all trials as operating under the high-reward conditions and fit one set of parameters for all trials. This was our best fitting model, as such the best model from the Expected Value Computation strategy was one that included a subjective non-linear cost accrual and treated high and low-rewarded trials as the same (see Supplementary Methods: All Sampling Strategies).

#### Evidence Difference Heuristic Strategy

Our second model, Evidence Difference Heuristic (ED), was a heuristic that assumed participants tracked the continuous signed difference in evidence between the two categories towards a threshold (e.g., “I sample until one category has 4 more than the other”). This strategy suggests that participants approach information sampling insensitive to the number of images sampled and implies that the stopping boundary is stationary and constant across sampling (Baumann et al., 2020; Korn & Bach, 2018; Shah & Oppenheimer, 2008; Herbert A. Simon, 1955; Tversky & Edwards, 1966). To fit this heuristic, we adapted the rule into a probabilistic account that used the signed difference in evidence drawn to predict choice. The signed difference in evidence between the current majority and minority in the samples collected at each timepoint was submitted to a multinomial SoftMax regression along with a subject-specific intercept, *β*_0_, in order to produce a probability for each action (Eq. 10).

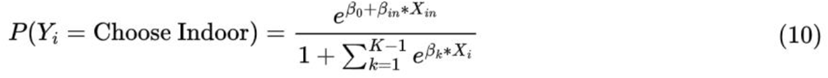

#### Sample Number Heuristic Strategy

Our third model, Sample Number Heuristic (SN), was another heuristic that assumed, participants used the continuous number of samples drawn and only a binary representation of the difference in evidence to inform choice (e.g. “I sample 5 images and then choose the majority). Similar to the first, this strategy maintained a stationary threshold that triggered a decision to select an option but implied that the magnitude of evidence mattered less. Identical to our Evidence Difference Heuristic Strategy, to fit this heuristic, we adapted the rule into a probabilistic account that used the number of samples drawn and a binarized difference in evidence to predict choice. These variables were submitted to a multinomial SoftMax regression along with a subject-specific intercept, *β*_0_, in order to produce a probability for each action (Eq. 11).

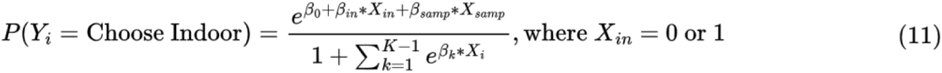

#### Evidence-by-Costs Tradeoff (ED-CA) Strategy

Our third series of models were built on the assumption that participants’ decisions to continue sampling or stop and commit to a category could be described by a strategy that depended on multiple forms of information but did not require the computational complexity of optimal strategies. Specifically, the Evidence-by-Costs Tradeoff (ED-CA) strategy, was a combination of the above heuristic strategies, such that participants used both continuous representations of the difference in evidence between the categories and the number of images collected to inform their choices. This strategy reflected a linear threshold that decreased as the number of samples collected increased, representing a non-stationary but constant tradeoff between the two informational sources. To fit this model, both sample number and the signed difference in evidence were submitted to a multinomial SoftMax regression along with a subject-specific intercept, *β*_0_, in order to produce a probability for each action (Eq. 12).

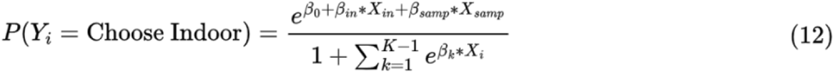

Similar to the Expected Value Computation model, we were also interested in whether participants treated information similarly across the different reward conditions. To test if reward significantly changed model fits, we tested two different model iterations. First, to test if participants were using completely difference parameter estimates for the different reward conditions, we split participant trials into high- and low-reward trials and fit each subset of trials to our SoftMax multinomial regression (Eq. 12). We then compared the difference in parameter values for high-vs. low-rewards. Second, to test if reward independently modified choice but did not impact the weight of individual information quantities, we added an additional reward parameter into the original SoftMax multinomial regression. Interestingly, parameter values were comparable across the two reward conditions and adding reward as an independent parameter did not improve model fit beyond Eq. 12. Thus, the best fitting model from the ED-CA Tradeoff strategy was one that also treated high and low-rewarded trials as the same (see Supplementary Methods: All Sampling Strategies).

#### Model Comparison

For each model, we optimized the parameters to maximize the log likelihood for each participant individually. We used SciPy’s standard optimize. minimize function to minimize the negative loglikelihood of the observed choices. Parameters for our Optimal Stopping were bounded based on previous studies (*p*: [0-25] for sigmoidal, *p*: [0, 0.2] for linear, *β*: [1,10]: [0, 0.5]) (Hauser et al., 2018) and both our Heuristic and Evidence-by-Cost models were bounded based on preliminary mixed effects multinomial regression [*β*_1_ (samples drawn): [−1,5], *β*_2_ (evidence difference): [−4,8], *β*_3_ (reward context): [−5,8]). In every case, we ensured the best fitting parameters each fell within these boundaries. We fit each participant 10 times per model to ensure convergence and stability of best fitting parameters.

To compare participants’ fits from our models, we first took the top performing models from each strategy if a strategy had more than one iteration before examining cross-group comparisons. All models in the final group were compared using both Akaike Information Criterion (AIC) (Akaike, 1974) and Bayesian Information Criterion (BIC) (Schwarz, 1978) scores. To examine patterns of best model fits on the group level we ran a repeated measures ANOVA to determine if participant-specific AIC or BIC scores differed significantly amongst models. Distributions of AIC and BIC scores per top performing can be found in the Supplement (see Supplementary Note 3, Figure 2).

## Statistics

All other statistics are stated in the text and figure captions. Normality was not directly tested because of our large sample sizes, but unless otherwise noted, data were assumed to be normally distributed and individual data points are provided in the figure scatterplots.

## Programming environments

Python 3 was used to run information sampling computation models and make data plots and figures. R, version 4.0.5, was used to calculate statistics (R Core Team, 2017).

## Code availability

Requests for the data can be sent via email to the corresponding author.

## Data availability

Requests for the code used for all analyses can be sent via email to the corresponding author.

