## Supplemental Materials for "Between heuristics and optimality: Flexible integration of cost and evidence during information sampling"

**Supplementary Information**

### Supplementary Methods: Task Instructions

In the first part of this study, we are interested in how collecting information can help guide your choices. Your task will be to collect information from a series of boxes. On each trial, the box will contain 25 images that can be categorized into two groups (e.g. indoor or outdoor scenes). In each box, there will be more images from one of the categories. For example, a box might have more outdoor pictures than indoor pictures. Your job on each trial is to figure out which category of images outnumbers the other category inside the box. You can collect images from the box one-by-one until you feel confident in your decision, at which point you will select the category that you think is most common in the box. You will then receive feedback about whether or not your choice matched the actual contents of the box. The exact proportions of images in the box may vary across trials but there will always be a correct category for each trial.

Each trial will also involve a potential reward for getting the answer correct. How much a trial is worth will be shown at the beginning of each trial on the "ready screen". Trials can be worth either $1.00 or $5.00. This "ready screen" will tell you what categories of images you will see in the trial and how much a correct response is worth for that trial. After the ready screen, a "sample screen" will appear where you will choose to either sample images from the box (center) or select your final answer for the trial (left or right side of the screen). If you choose to sample, an image will appear on the screen for a couple seconds. After the picture disappears, you will be returned to the "sample screen" where you can decide if you want to sample from the box again or make your final choice. The images inside the box have been shuffled so the pictures you see in the beginning may or may not reflect the rest of the box's contents.

There is also a cost for sampling images., Each time you choose to sample from the box, a small deduction will be made from your potential total earnings for that trial. For example, for $5.00 trials, the cost to sample is $0.10; thus, every sampled picture subtracts 10 cents from your total potential reward for that trial. For $1.00 trials, the cost to sample from the box is $0.02. A correct response means that trial is worth the original reward minus sampling cost (e.g. sampling 7 times makes a correct response worth $4.30), an incorrect response means a trial is worth $0.00. The values of these trials are what is used to calculate your bonus.

You can sample up to 25 images from each box. If you have not yet made a choice after the 25th sample, you will be prompted to choose a category. When you feel confident that you know which category out numbers the other category in the box, select that category. You will receive feedback about whether or not your choice was correct after your final decision on a "feedback screen." You will then move on to the next trial, where you will sample images from a new box.

Throughout the study, you will use the three arrow keys at the bottom of the keyboard to make your responses. These arrow keys will line up with the options on the screen such that the left arrow means you are choosing the category on the left side of the screen, the down arrow means you are choosing to sample, and the right arrow means you are choosing the category on the right side of the screen. At the end of the experiment, you will randomly select 3 trials from the experiment. Your bonus will be the sum of the earnings from these three trials. This means that your bonus can be calculated from trials that were associated with either high or low reward and from trials that were answered either correctly or incorrectly. Therefore, it is best to try and get every trial correct because each could contribute to your final bonus.

Do you have any questions? Ready to practice?

#### Supplementary Note 1, Figure 1: Participant total earnings correlated with realized bonus

Participants completed 48 trials of the information sampling task. Half of the trials were completed under low monetary stakes ($1.00 potential reward) and half were completed under high monetary stakes ($5.00 potential reward). Participants received a performance bonus that depended on a subset (4, randomly selected) of the trials of the 48 completed trials. Participants accumulated an average of $95.07 across all trials (SD = $10.52). The average realized bonus was $9.25 (SD = $3.05). The total amount participants earned across the entire task was positively correlated with their realized bonus (Pearson's $r(92) = 0.31$, $p = 0.002$).

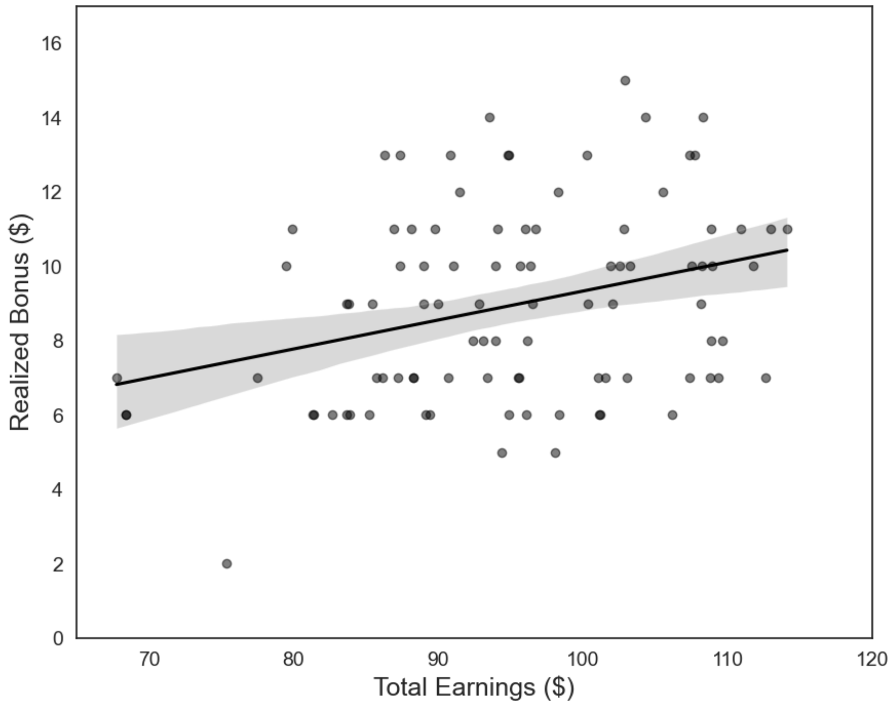

Supplementary Figure 1: Participant total earnings correlated with realized bonus. The total amount of money accumulated during the sampling task positively related with the actual performance bonus participants received. Each dot represents one participant.

### Supplementary Methods: Ideal Observer Model

To assess performance for an ideal observer, we first constructed a model of optimal behavior. We defined optimality as a policy that maximized the expected value of each choice across each trial. Expected value was computed as the probability a correct choice would be made, and as such reward would be obtained, minus the costs accrued from sampling. Thus, similar to our human participants, behavior optimized to maximize reward on each trial, needed to balance the probability of successfully identifying the underlying majority category with the costs of collecting information. Our model of optimal behavior utilized similar computations as the Expected Value Computation strategy (see Supplementary Methods: Expected Value Model Formulation), such that the ideal observer computed the value for taking all available actions (continue sampling, choose indoor, or choose outdoor) and took the highest valued action. Unlike the probabilistic model described below, costs were integrated objectively, and choices did not include noise. We ran our ideal observer through our task 94 times using the same number of trials as our participants completed. We then computed the average accuracy across all runs as well as the average total earnings across all trials.

### Supplementary Methods: Expected Value Model Formulation

The specific set-up for information sampling in this task is as follows. On a given trial, participants are told that there is a box containing 25 images within. Each image belongs to one of two categories (e.g., indoor vs. outdoor scenes) and the proportion of images from one category against the other is unbalanced. Participants are given the goal of identifying which category holds the underlying majority for that box. They can do so by sequentially sampling images from the box until they feel confident enough to decide. Correct decisions are rewarded and incorrect decisions result in no reward as well as no loss in reward. Each image sampled comes with a cost that is deducted from their potential reward.

The specific information provided to a participant for any given trial is:

- The box contains a total of 25 unique images
- The maximum reward value for a trial is either $5.00 or $1.00
- The cost per image is a constant 2% of the maximum reward available on a trial ($0.10 for $5.00, $0.02 for $1.00)
- The proportion of images from either category is specifically withheld and participants are told that the proportion may change on a trial-by-trial basis

The current paper explores several potential strategies that could guide information sampling decisions. One potential strategy involves estimating the expected value of each available action through inferring the true state of a box using Bayesian reasoning. Below is a detailed account of the formulation for such model.

***Estimating the Probability of Success Given Available Evidence.***

To estimate the expected value of each action, one must first compute the probability of that choosing a particular category given the available evidence will result in the correct identification of the majority. Participants know at any given point what their current evidence is, that is how many images they have sampled and how many of those images belong to the a specific category (e.g. indoor scenes). To start, we will set up the logic for estimating whether the underlying majority is Indoor given our current evidence as such:

$$P(Indoor|n_i, N)$$

Because participants are aware that there are only 25 images in the box, they know that once 13 images from one category have been sampled, there is certainty about the majority category:

$$P(Indoor|n_i, N) = \begin{cases} 1 & \text{if}\, n_i > \frac{N_{tot}}{2} \\ 0 & \text{if}\, N - n_i > \frac{N_{tot}}{2} \\ & \text{else, next eq.} \end{cases}$$

When there are less than 13 images from one category in the current evidence, the probability that a box contains a majority of indoor images must be inferred by the participant. To make this inference, several steps are to take place. First, since the underlying number of indoor images is unknown to the participant, the participant must estimate what the underlying distribution of indoor to outdoor scenes is in order to then estimate the probability the majority of images are indoor scenes. Estimating the likelihood of any combination of indoor to outdoor images is broken up into two computations. First, we estimate the underlying generative probability (q), given the currently available evidence. The generative probability can be any probability between 0 and 1, and is integrated over all possible values of q.

$$1 = \int_0^1 P(q|n_i, N)dq$$

We estimate the underlying generative probability for each potential combination of indoor and outdoor images left in the box. Thus, we are inferring a weighted average of what remains in the box given the evidence we have currently gathered.

$$P(Indoor|n_i, N) = \sum_{Indoor>\frac{N_{tot}}{2}}^{N_{tot}-(N-n_i)} \int_0^1 P(Ind|q, n_i, N)P(q|n_i, N)dq$$

The above equation can be further delineated to reflect the estimation only of the evidence yet to be sampled, that is, the total images minus the ones already drawn.

$$P(Indoor|n_i, N) = \sum_{Indoor>\frac{N_{tot}}{2}}^{N_{tot}-(N-n_i)} \int_0^1 P(Ind - n_i, N_{tot} - N|q)P(q|n_i, N)dq$$

The first part of the equation, \$P(Ind-n_i,N_{tot}-N|q,n_i, N)\$ is a binomial distribution, and as such, can take the form:

$$Bin(Ind - n_i|q, N_{tot} - N)$$

Binomial distributions are described by two components: a combinatorial of all possible arrangements of pulling k successes in n draws (or k indoor images out of n images drawn) and the likelihood of successes and failures given an underlying probability.

$$\binom{n}{k} p^k (1-p)^{n-k}$$

Thus, this distribution can be expanded to our task demands such that, we are determining the combinatorial for how many indoor scenes could be drawn from the remaining images still in the box and estimating the likelihood of successes and failures (drawing indoor vs. outdoor images) using the currently gathered evidence as such:

$$\binom{N_{tot} - N}{Ind - n_i} q^{Ind-n_i}(1-q)^{N_{tot}-N-Ind+n_i}$$

The second part of the equation, $P(q|n_i, N)dq$ is a beta distribution, which has a PDF of:

$$\frac{x^{\alpha-1}(1-x)^{\beta-1}}{B(\alpha,\beta)}$$

With $\alpha, \beta$ are shape parameters. We set a flat prior on q, $\alpha = \beta = 1$, which means our prior essentially equals 1 (and we won't really mention it further). If we adapt the beta distribution to include the current evidence gathered, we can expand our estimation to:

$$P(q|n_i, N) = \frac{q^{\alpha+n_i-1}(1-q)^{\beta+(N-n_i)-1}}{B(\alpha+n_i, \beta+N-n_i)}$$

To break this down further, the first element of the numerator $q^{\alpha + n_i - 1}$ reflects the number of successes (indoor images) plus the original beta function of $\alpha -1$. Similarly, the second element of the numerator, $(1-q)^{\beta + (N-n_i)-1}$, reflects the number of failures (non-indoor images) plus the original beta function of $\beta - 1$. Now, combining like terms, we can calculate:

$$P(Ind - n_i|n_i, N) = \int_0^1 P(Ind - n_i|q, N_{tot} - N)P(q|n_i, N)dq$$

$$= \frac{\binom{N_{tot}-N}{Ind-n_i}}{B(\alpha+n_i, \beta+N-n_i)} \int_0^1 q^{Ind+\alpha-1}(1-q)^{N_{tot}-Ind+\beta-1}dq$$

$$= \binom{N_{tot} - N}{Ind - n_i} \frac{B(Ind+\alpha, N_{tot} - Ind + \beta)}{B(n_i+\alpha, N-n_i+\beta)}$$

The last component is to add back in the summation. We are computing this probability over all possible combinations of majority Indoor images, so the final form looks like:

$$\sum_{Indoor>\frac{N_{tot}}{2}}^{N_{tot}-(N-n_i)} \binom{N_{tot}-N}{Ind-n_i} \frac{B(Y+\alpha, N_{tot}-Y+\beta)}{B(\alpha+n_i, \beta+N-n_i)}$$

Importantly, this computation reflects the probability that the underlying majority category is indoor. The same computation can be performed with $n_o$ instead of $n_i$ to get the probability that the underlying majority category is outdoor. Both computations are necessary to determine how likely a particular choice with happen at any given time point.

***Constructing Action Values for Choosing Indoor vs. Outdoor***

Now that we have a single estimate for the probability of successfully identifying the underlying majority as Indoor (and Outdoor), we can use those probabilities to compute the action value for actually choosing the category Indoor (or Outdoor). To do so, we multiply the Reward for correct identification with the probability of success for each choice:

$$Q(Indoor|n_i, N) = R_{correct} P(Indoor|n_i, N)$$

$$Q(Outdoor|n_o, N) = R_{correct} P(Outdoor|n_o, N)$$

For this task, the reward for correct identification is either \$5.00 or \$1.00. The action value for each stopping choice does not contain information about the value of choosing the other category because there is no penalty for incorrectly identifying the majority. Also note, that in this version of this model, the accumulated costs are not represented in the above stopping values. The reason for this is to consolidate both the objective costs of sampling and the subjective costs of sampling into a single estimate.

***Constructing Action Value for Continuing to Draw Samples***

To evaluate decision options, the expected values of the choices to stop and select a majority category must be compared to the expected value of the choice to continue drawing images. In general, we start with the notion that the value of continuing to draw images can be computed by taking the expected values of future states, weighted by the likelihood of reaching those future states minus the cost of continuing to sample. Thus, the formula for computing the action value of continuing can be written as such:

$$Q(Continue|n_i, N) = -c_{\text{per step}} + \sum_{\text{all possible s'}} P(s'|n_i, N)V(s')$$

Because the value of continuing involves looking into the future and estimating potential trajectories of choices, this computation involves several steps:

1. $V(s')$: The value of a future state is computed. This involves:
    1. Determining a global policy ($\pi$) that maps the probabilities of taking particular actions, a, in particular states, s (or s').
    2. Using that policy, taking note of the actions available at state s' and computing an action value for each action.
    3. Summing across all available actions in s' weighted by the likelihood that action is chosen (from $\pi(a,s')$) to give a single value estimate, $V(s')$
2. $P(s'|n_i,N)$: The probability of reaching a particular state, s', is computed. This involves:
    1. Computing the probability that the next image pulled will be indoor (or outdoor) given the current evidence and integrating over q
3. $-c_{\text{per step}}$: The cost of taking a step into the future is computed and added to the equation.

The equation above has two summation components to it. One summation component is part of the construction of $V(s')$, where we are taking the action value of each available action at s' (usually 3—choose outdoor, choose indoor, choose draw sample—or 2 if at the last step), multiplying it by the probability that that action is chosen (derived from the policy, $\pi$), and then summing across all action by probability pairings. The second summation happens when we take the average of the two $V(s')$'s. At most timepoints, it is possible to move forward in two directions (N + 1, ni + [0 or 1]), therefore, there are two potential states that can be reached from s. Since we do not know what category the next image will come from, we estimate the probability of getting to each individual s' and then multiple that transition probability with the value of that state, $V(s')$. We do this for each s' and then sum them together to get a single estimated value for continuing to sample information. All of these steps are wrapped into a general function that performs backward induction to derive the values of all potential states based on the value of the last possible state that can be reached.

*The value of a future state.* Generally, this computation can be thought of as taking the expected value of the potential next states (steps into the future, s') that will then be weighted by the probability of actually reaching those states (next function below).

$$V(s') = \sum_{\text{all available a in s'}} \pi(a, s')Q(a, s')$$

To find the expected value of the potential next state, we first assume a particular policy a participant is using.

$$\pi(a, s')$$

This policy gives us the probabilities that particular actions will be taken at s'. To estimate these probabilities, we assume that participants are using a Softmax function across the three available choices (Continue sampling, Stop choose Indoor, Stop choose Outdoor).

$$\pi(Choice|Evidence) = \frac{\text{Each Choice individually}}{\text{All Choices}}$$

This SoftMax is then individualized for each participant with the parameters $\beta$ (inverse temperature) and $\xi$ (irreducible noise):

$$\pi(Choice|n_i, N) = \frac{e^{Q(Cont.|n_i,N)\beta}}{e^{Q(Cont.e|n_i,N)\beta} + e^{Q(Ind.|n_i,N)\beta} + e^{Q(Out.|n_i,N)\beta}}(1-\xi) + \frac{\xi}{3}$$

Once we have our policy (which again computes the probability that a particular action will be taken), we then multiple each probability by the action value for that particular action as reflected in:

$$\sum_{\text{all available a in s'}} \pi(a, s')Q(a, s')$$

Each s' (besides at time point 25), we will have three probabilities from our policy function and three action values corresponding to the available actions at s' (namely, continue, stop-indoor, stop-outdoor). Each probability and action value are multiplied together and then summed over all available actions in that s' to get $V(s')$. Importantly, the value of s' differs by whether the next sample drawn is another indoor image or outdoor image. This means that $V(s')$ is computed twice, once when the next sample drawn is indoor and one when it is outdoor.

$$Q(Continue|n_i, N) = -c_{\text{per step}} + \sum_{s'=\begin{cases} n_i + x \\ N+1 \end{cases}}^{x=[0,1]} P(s'|n_i, N)V(s')$$

These individual computations are then summed together. We will return to the calculation of these probabilities in the policy and the calculations of the action values in s' when we discuss backward induction.

*Probability of reaching a future state.* After computing the value of taking a step into the future, those values are multiplied by the probability of reaching that particular state, given the current evidence collected.

$$P(s'|n_i, N)$$

For this task, this is the probability of the transitioning from $N, n_i$ → $N + 1, n_i + 1$ or $N+1, n_i + 0$.

$$= P(n_i + x, N + 1 | n_i, N)$$

Where x can equal to 0 or 1. To compute the transition probabilities, we must again estimate and integrate over the value of q as such:

$$= \int_{\text{all q}} P(x|q)P(q|n_i, N)dq$$

These can be broken up in the a Binomial distribution, where the value of x depends on the value of q, and a Beta distribution to estimate the value of q (the probability that q is q integrated over all possible values of q).

$$= \int_{\text{all q}} Bin(x|1, q)P(q|n_i, N)dq$$

Breaking down the Binomial distribution, as before, we have both a combinatorial element (n choose k) and a product reflecting the probability of drawing k and n - k amount of successes and failures. Because we are examining one step into the future, the combinatorial here is either equal to:

$$\binom{1}{0} \text{ or } \binom{1}{1}$$

both of which equate to 1. Breaking apart the beta distribution and adding in the components from the Binomial distribution like before, this can be rewritten like:

$$\frac{B(\alpha + n_i + x, \beta + N - n_i + 1 - x)}{B(\alpha + n_i, \beta + N - n_i)}$$

The numerator is a beta function with the two shape parameters, that are adjusted to include the additional step and resultant evidence that comes from that step. The $\alpha +n_i+x$ is the immediate future step with the "successes" (aka number of indoor images) altered by x (so either + 1 or + 0). The $\beta + N - n_i + 1 - x$ is the immediate future step with the number of "failures" altered by + 1 - x (again either 0 or 1 but reversed from the number of successes). The normalizing term is the same as previously used.

*The cost per step.* Lastly, the cost of sampling per step, $c_{\text{per step}}$, is subtracted from the value of continuing as shown above. In this iteration of the model, the cost has a sigmoidal form, suggesting that participants are insensitive to the accumulating costs until a particular sample number is reached, at which point costs scale up to the objective cost per sample.

$$c_{\text{per step}} = \frac{\text{objective cost}}{1 + e^{-k(n-p)}}$$

Here, the numerator reflects the objective cost per step for sampling an image and the upper boundary of $c_{\text{per step}}$. $-k$ reflects the slope of the sigmoid (here set to 10), $n$ reflects the number of samples drawn and $p$ reflects the change point at which costs start accumulating (also referred to as the "patience" parameter).

*Backward Induction to derive action values and action probabilities.* Since the horizon in finite (there are only 25 images that can be drawn), the value of continuing at the end of 25 images is 0 and the value of stopping is fully determined (Probability of Success = 1). Knowing this, we can work backwards to determine the value of the available actions at time point 24, 23, 22, etc. We start with time point T = 25, which reflects drawing all available images from the box. At T= 25, the value of continuing to sample is equal to 0 since there are not more images to be sampled. The value of stopping and choosing a majority category is equal to:

$$V_{\text{Indoor, T = 25}} = Reward * P(Indoor)$$

The $P(Indoor)$ and the $P(Outdoor)$ will be either 1, 0 or 0, 1 depending on the underlying majority, which will have been realized given that there are no more images in the box. From here, we can move backwards to time point t = 24 and compute the value of continuing to T = 25 and compare it to the value of stopping at t = 24. Computing the value of continuing relies on the steps listed in *Constructing Action Value for Continuing to draw samples* and computing the value of stopping relies on the computations listed in *Constructing Action Values for Choosing Indoor vs. Outdoor***.**

We calculate the value of continuing and stopping (one value for each stop option) across all potential combinations of observed evidence (e.g., at t = 24, N = 24, Y = 24; N = 24, Y = 23; N = 24, Y = 22;...; N=24, Y = 0). At the end, we have a table that holds for each possible step, the expected value of that position, which reflects the expected values of all potential steps to come. The table also holds the probabilities determined by $\pi$ of the probability of choosing each action at state s. A key element is there is an assumption in backwards induction that the policy is the same in the creation of the table of values and the execution of moving through those values (policy consistency).

***Deciding Amongst Actions Given the Current Evidence***

To understand how participants are selecting between the various actions available at any given time point, the action value for each choice is computed using the action value for stopping choices, or the inverse for the action value for choosing outdoor, and the EV's derived from the lookup table for s' multiplied by the transition probabilities.

$$Q(Indoor|n_i, N) = R_{correct}P(Indoor|n_i, N)$$

$$Q(Continue|n_i, N) = -c_{\text{per step}} + \sum_{s'=\begin{cases} n_i+x \\ N+1 \end{cases}}^{x=[0,1]} P(s'|n_i, N)V(s')$$

The same policy function is then used to derive the probabilities associated with the choice options, where the numerator changes to get the probability of each choice.

$$\pi(Choice|n_i, N) = \frac{e^{Q(Cont.|n_i,N)\beta}}{e^{Q(Cont.e|n_i,N)\beta} + e^{Q(Ind.|n_i,N)\beta} + e^{Q(Out.|n_i,N)\beta}}(1-\xi) + \frac{\xi}{3}$$

***Estimating a Participant's Free Parameters***

To find the best estimate of a participant's parameters given their actual choices, several steps are performed. First, we initialize our free parameters randomly within the pre-determined range of the parameter space (See main text). We then compute the negative log-likelihood across all choices made given a specific set of parameters. To do this, we take the probabilities computed from the Softmax function above, see which choice a participant actually made at any given time point and then take the log of that probability associated with the actual choice made. This is done for each choice and the log values are stored in an array. We then take the negative sum over the array of log likelihoods. We use a maximum likelihood estimator to iterate over different parameter values until we find the combination of parameter values that result in the largest negative log-likelihood (moving as close to zero as possible). These values represent the free parameters that best describe the participant's behavior.

### Supplementary Methods: Beta-Binomial equivalent to Hypergeometric when K in unknown

We use the Beta-Binomial distribution to estimate the probability of success given the current evidence. Although our task technically drew from a Hypergeometric distribution, we show here that when K is unknown, these distributions produce equivalent results.

First, let us assume a hypergeometric distribution in which the urn contains N balls and K total successes. Then the probability of observing k successes in n draws is

$$P(k|n,K,N) = \frac{\binom{K}{k}\binom{N-K}{n-k}}{\binom{N}{n}}$$

We further assume a Beta-binomial prior on K given N:

$$P(K|N,\alpha,\beta) = \binom{N}{K}\frac{B(K+\alpha, N-K+\beta)}{B(\alpha,\beta)}$$

It is tedious but not hard to show that, under these assumptions, the posterior is given by

$$P(K|k,n,N) = \mathrm{BeBin}(K-k, N-n, \alpha+k, \beta+n-k)$$

$$= \binom{N-n}{K-k}\frac{B(K+\alpha, N-K+\beta)}{B(\alpha+k, \beta+n-k)}$$

which is equivalent to the expression above under the substitutions

$$K \to Y = Ind$$
$$k \to n_i$$
$$N \to N_{tot}$$
$$n \to N$$

That is, the assumption of a binomial model on future draws (as calculated previously) is *equivalent* to the assumption of a hypergeometric distribution over draws when we use a Beta-binomial prior over the number of successes K in the urn.

### Supplementary Methods: All Sampling Strategies

We tested several variants of our top two strategies (Expected-Value Computation and ED-SN Linear Tradeoff) to explore the extent to which reward and cost altered information sampling. For the Expected-Value Computation, we first tested which of 3 potential cost functions best described how participants were integrating the cost of drawing an image. In the sigmoidal cost function, costs accumulated as a single step function, such that costs did not accumulate until a certain number of samples had been drawn, at which point costs start to accumulate at the objective sampling cost rate of 2% of the max reward. The patience parameter ($p$) represents the number of samples drawn before the costs start to accumulate. In the linear cost function, costs accumulate linearly and are scaled by how sensitive participants generally are to the sampling costs. Here, the scale parameter ($t$) represents the linearly scaled version of the objective costs. Our last cost function determined sampling costs as an ideal observer would, as such, this model had no additional free parameter. Of these models, the sigmoidal cost function best fit our data, replicating results for Hauser et al. 2018.

Across all these variants, we collapsed the high and low reward trials together, such that all trials were run using the expected values generated from the high reward condition. This was done to provide greater separation of action values before submitting them to the SoftMax function. We then examined whether participants were using difference strategies for our different reward conditions. We adapted the Expected-Value Computation with the sigmoidal cost function into 2 additional variants. One variant generated the expected values of each reward condition using the maximum reward for that trial, but low-reward trials could be scaled up ($s$) to a maximum of $5.00. Our other variant fit separate parameters for high- and low-reward conditions and included a scaling parameter for low-reward trials. Neither variant outperformed the original Expected-Value Computation with the sigmoidal cost function.

In our ED-SN Linear Tradeoff variants, we tested three models in total. Our first model tests the descriptive power of an intercept and both samples drawn and evidence difference. Our second model tests the descriptive power of an intercept, samples drawn, evidence difference, and the reward condition. Our last model generated separate parameters for the high- and low-reward trials. Across the three tested models, the best fitting variant was one that included parameters for an intercept, samples drawn, and evidence difference. Including reward information did not improve fit nor did fitting separate parameters for the different reward conditions.

Supplementary Table 1 indicates the model name, number of parameters, negative log-likelihood, AIC, and BIC scores for all tested models. Negative log-likelihood, AIC, and BIC are summed across all participants included in the analyses. We used BIC to determine the best model performance because of the stricter criteria for each additional parameter.

| Model Name | # of Parameters | negLL | AIC | BIC |
|---|---|---|---|---|
| EVC with Sigmoidal Cost Accumulation | 3 | 10192 | 20856 | 21981 |
| EVC with Linear Cost Accumulation | 3 | 11348 | 23170 | 24295 |
| EVC with Objective Cost Accumulation | 2 | 13161 | 26608 | 27358 |
| EVC with Sigmoidal cost and low-reward scale | 4 | 10132 | 20914 | 22414 |
| EVC with sigmoidal cost, separate parameters by reward | 7 | 9998 | 21214 | 23839 |
| SN-Heuristic | 3 | 11355 | 23184 | 24309 |
| ED-Heuristic | 2 | 10572 | 21416 | 22166 |
| ED-SN Linear Tradeoff | 3 | 8742 | 17964 | 19089 |
| ED-SN Linear Tradeoff + Reward | 4 | 8577 | 17818 | 19318 |
| ED-SN Linear Tradeoff, separate parameters by reward | 8 | 8227 | 17870 | 20871 |

Supplementary Table 1. Comparison of fits across tested models of behavior. Group negative log-likelihood, AIC, and BIC scores were calculated by summing individual scores from each participant for each model. An equal number of participants contributed to all model scores.

### Supplementary Methods: Parameter Recovery and Model Recovery

To examine how well each model could recover parameters from data generated from that model, we created 94 fake participants that experienced identical task conditions experienced by our participants. Each instance of generated data completed 48 trials, 24 in the high reward and 24 in the low reward conditions. The sequence of indoor and outdoor images was randomized identically to the real experiment. Data from each generation was submitted to the same fitting as our participants. Plotted below are the differences between the true and estimated parameter values for each parameter for each model.

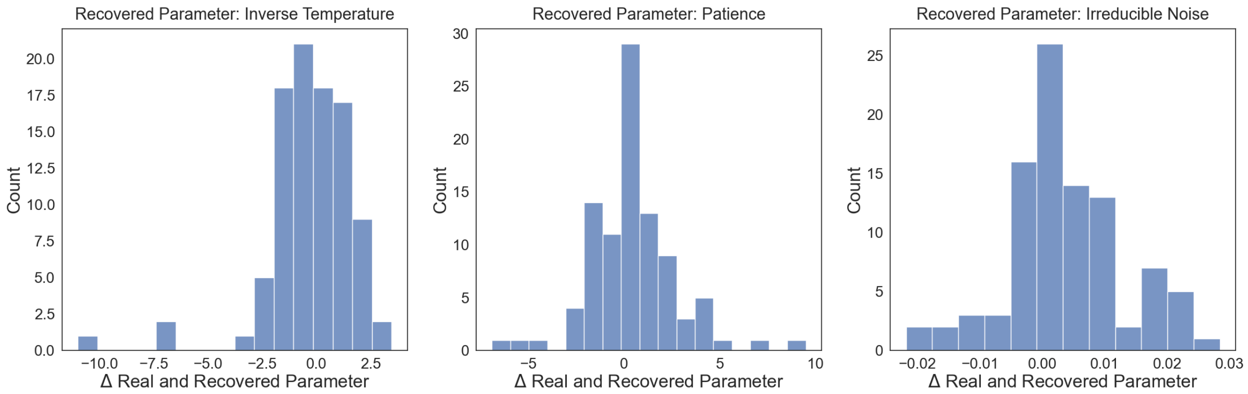

Supplementary Figure 2: Parameter recovery for winning Expected Value model. A) Difference between true and recovered parameter for inverse temperature parameter, Beta, B) difference between true and recovered parameter for patience parameter, p, C) difference between true and recovered parameter for irreducible noise parameter, xi. Data generated under the same task conditions participants experienced.

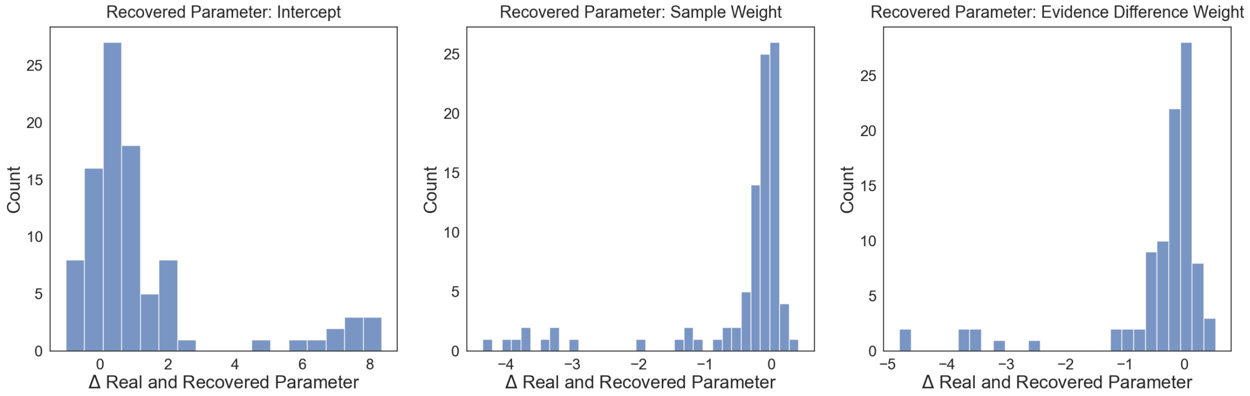

Supplementary Figure 3: Parameter recovery for winning Linear Threshold model. A) Difference between true and recovered parameter for fitted intercept, Beta_0, B) difference between true and recovered parameter for fitted weight on sample number, Beta_1, C) difference between true and recovered parameter for fitted weight on evidence difference, Beta_2. Data generated under the same task conditions participants experienced.

In addition to testing parameter recovery, we also checked whether data generated by one model was best fit by the model that generated it. To test this, we took our 94 simulations of generated data from each model and fit each instance to each of our winning models. For each simulation, model parameters were sampled randomly for the within the parameter bounds specified by each model. Each simulated data set was then fit to each of the given models to determine which model fit best, according to BIC scores. Results demonstrated that of the 94 instances generated by the Expected Value models, 88 of them were best fit by the Expected Value model. Similarly, of the 94 instances generated by the Linear Threshold model, 90 of them were best fit by the Linear Threshold model. Thus, data generated by each model is best fit by the model that generated the data.

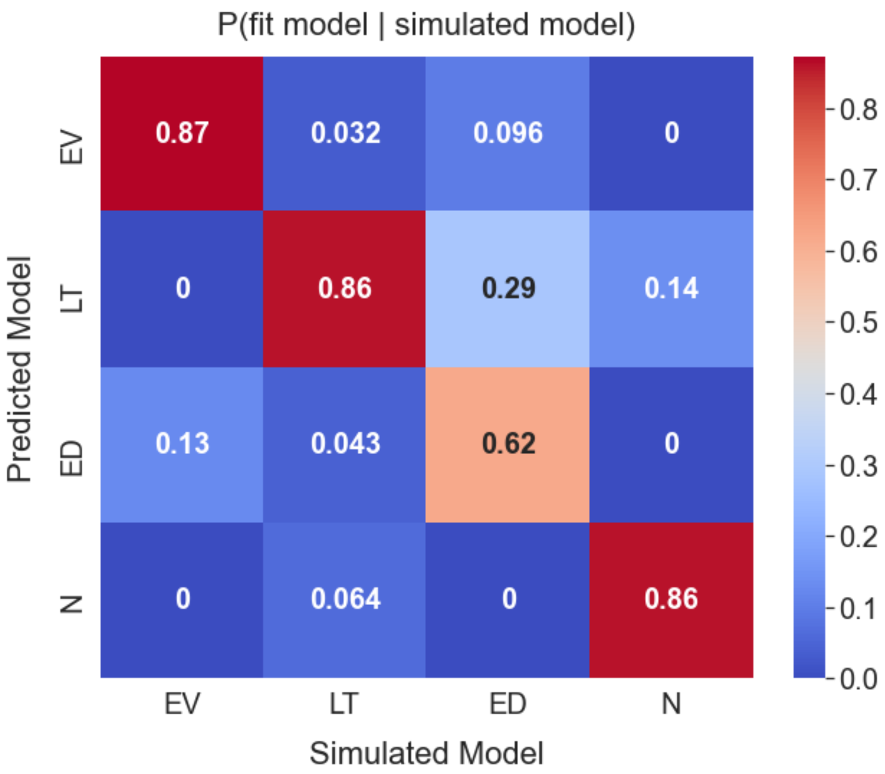

Supplementary Figure 4: Confusion matrix for top Linear Threshold model and Expected Value model. Each model was simulated 94 times and fit to both models. Best model fits were determined using BIC.

### Supplementary Note 3, Figure 2: Distribution of BIC scores by model.

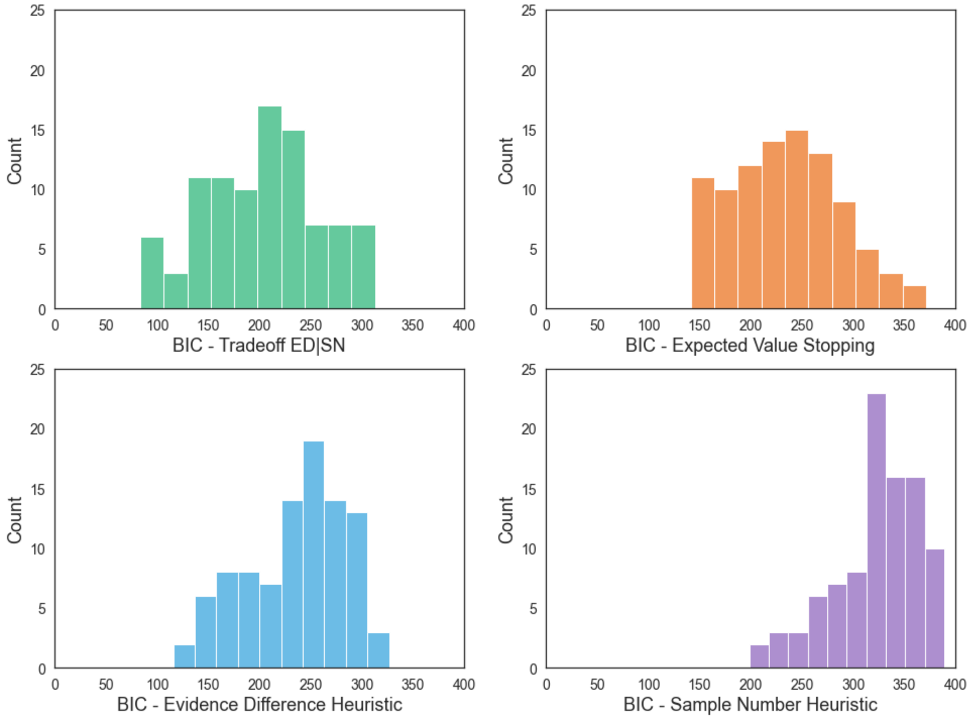

Figure 2. Distribution of BIC (top) and AIC (bottom) scores across the top-performing model for each family.

#### Supplementary Note 4, Figure 3: Reward motivation does not alter underlying stopping model

To investigate whether reward condition changed the underlying process participants used to arbitrate between sampling and stopping, we tested the impact of reward condition on model fit across the two best fitting models from each family. For each participant, task data was split into high and low reward trials and run separately through the Optimal-Based Stopping model and the Hybrid Stopping model. Additionally, because we split by reward condition, we removed the reward parameter for our Hybrid Stopping model. We fit each participant 10 times per model, per reward condition to ensure convergence and stability of best fitting parameters. Parameter boundaries were identical to those used in main analyses. BIC scores were used to assess the best fitting model across reward conditions.

Across participants, the Hybrid Stopping model best fit behavior in both high and low stakes trials. 19 participants had inconsistent best model fits between reward conditions. Of those participants, there was no consistent direction of change: Optimal-Based Stopping provided a better fit under high stakes trials for 7 participants and provided a better fit under low stakes trials for 12 participants. Thus, reward motivation did not reliably or consistently change the process model that best described participants behavior.

<table>
<tr><td colspan="2"></td><td colspan="2">High Stakes Trials</td></tr>
<tr><td colspan="2"></td><td>Hybrid Stopping</td><td>Optimal-Based Stopping</td></tr>
<tr><td rowspan="2">Low Stakes Trials</td><td>Hybrid Stopping</td><td>69</td><td>7</td></tr>
<tr><td>Optimal-Based Stopping</td><td>12</td><td>6</td></tr>
</table>

Supplementary Table 2. Increasing reward stakes does not increase the use of a strategy more akin to Optimal-Based Stopping. Participant task trials were split into high and low stake trials. Separate modeling of both high and low trials revealed an overall consistency in Hybrid Stopping providing the best fit for participant behavior.

### Supplementary Methods: Memory Task, Description, and Findings

Of the 104 participants to complete the first session, 102 participants returned 24 hours to partake in a surprise memory test. From the 102 participants, 16 participants were excluded from all analyses, eight due to computer error and eight due to having unusable sampling data (failed to sample at least once on over 25% of trials), leaving a final total of 86 participants. In the memory test, participants saw images they saw during information sampling in addition to new images that were not seen during the first session. Since the number of images seen during the first session varied across participants, we varied the number of new images seen such that of the total images seen in the memory test, 75% were old images and 25% were new images. All old and new images were randomized and presented one at a time. Participants first judged in the image was old or new and then rated their confidence in the memory judgement (Supp. Fig. 2).

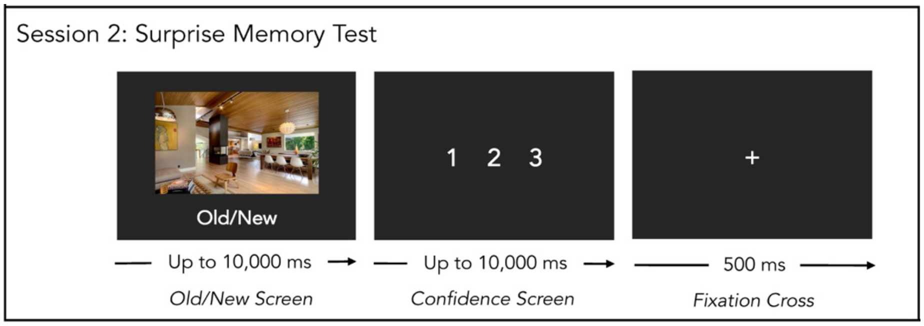

Supplementary Figure 2: 24 hours later, participants returned to the lab to complete a surprise memory test. Each image in the memory test was presented on the screen with the text "Old/New" presented below. Participants had up to 10 seconds to respond to each image. After they made an Old/New judgement, they were then given 10 seconds to rate their confidence in their memory judgement on a scale from 1 to 3, (1 = guess, 3 = sure). Participants then saw a fixation cross that separated each image.

Across participants, the average memory performance as 59% (SD = 7.2%). The mean number of old images presented was 374.8 (SD = 134.2) and the mean number of new images presented was 124.7 (SD = 44.7). Overall, participants appeared to understand the memory test and correctly used the confidence scale.

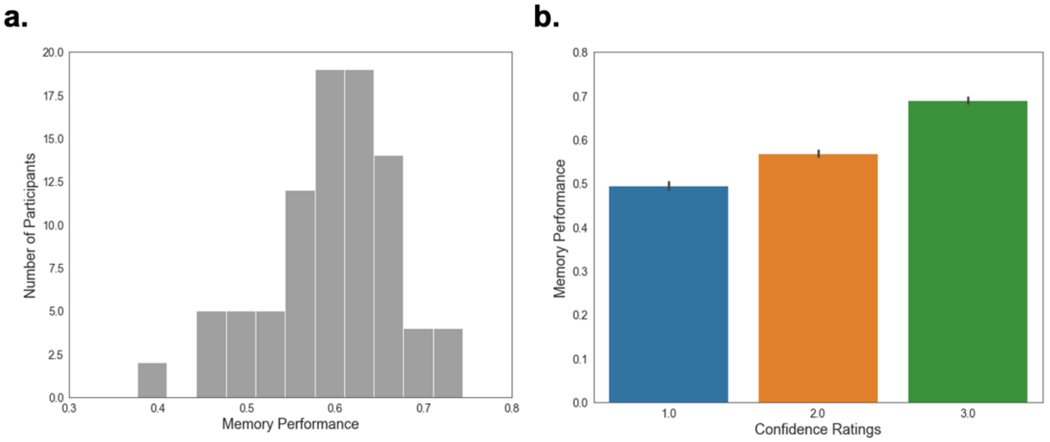

Supplementary Figure 3: Summary memory data for second session. a) Distribution of memory performance for second session. Memory performance for each participant was first computed and then plotted in the above history. Performance was marked as correctly identifying old images as one and new images as new regardless of confidence rating. b) Memory performance by confidence ratings. Higher ratings of confidence in memory judgement were associated with increased memory performance.

We first examined whether the total number of images a participant saw during the memory test impacted their memory performance. We first computed the corrected hit rate for each participant by subtracting the percentage of false alarms (new images that were judged as old) from the hit rate (old images judged as old) to get the corrected hit rate. We found that the total number of images seen during the second session did not significantly correlate with corrected hit rate (Pearson's $r(84) = -0.15$, $p = 0.163$). This suggests that memory performance was not significantly impacted by the load of images.

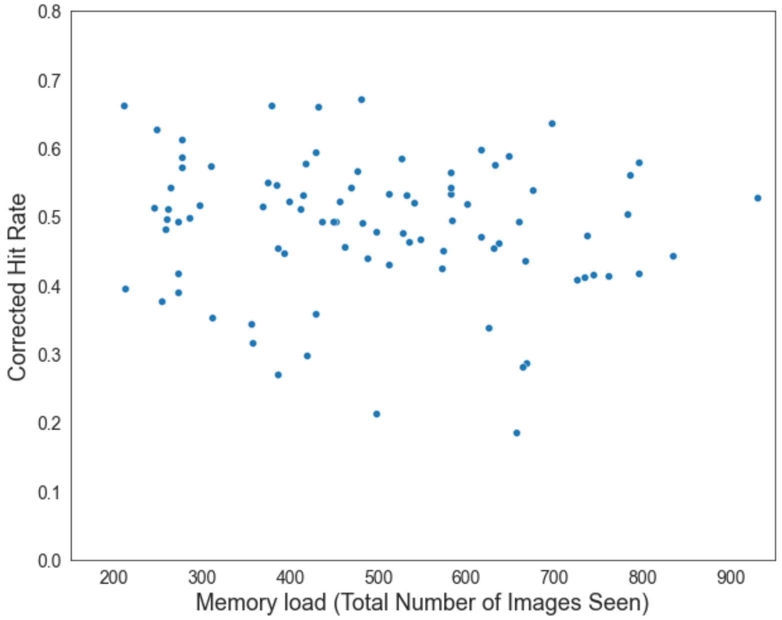

Supplementary Figure 4: Memory performance was not correlated with memory load. For each participant, the total number of images seen during the second session and the corrected hit rate for that subject was computed. Each dot reflects a single participant.

We tested four hypotheses about the potential relationship between sampling and memory formation. Our first hypothesis predicted that participants would preferentially remember images from high reward trials compared to low reward trials. Our second hypothesis predicted that participants would preferentially remember images from the category they choose over the one they didn't, and that this would be modulated by whether the choice was correct or incorrect. Our third hypothesis predicted that participants would preferentially remember images immediately prior to stopping decisions over information gathered early in the trial. Our last hypothesis predicted that participants would preferentially remember images when uncertainty was high compared to when it was low.

We conducted a single mixed effects logistic regression (Bates, Mächler, Bolker, & Walker, 2015) to test for support for any of our four hypotheses. Within the model, we included a covariate for the total memory load a participant experienced. Participants were treated as random intercepts. To our surprise, our model did not provide support for any of our hypotheses (all $p > 0.05$). At the current state, we cannot provide evidence elucidating if and how information sampling impacts memory formation for the sampling experience.

### Supplementary Methods: Response Time Analyses

Response times for both choices to stop or continue sampling that were longer that 5 seconds or were more than 2 standard deviations above each participant's mean were discarded and not analyzed further. The remaining values were Z-scored for each participant and aggregated across participants.

To investigate the relationship between response time and choice type (decisions to stop sampling vs. continue sampling), we tested whether choice type predicted response times in a mixed effects linear regression with subject as a random intercept (Eq. S3). We found that decisions to stop sampling were associated with lower response times compared to decisions to continue sampling (Stopping = -0.644, t = -23.4, p < 2e-16). This suggests that subjects were faster in making their final decisions compared to the decision to sample another image (Supplementary Figure 3a).

$$RT = \beta_1(choice\ type) + (1|subj) \qquad \text{(S3)}$$

We next examined whether this relationship was stable as a function of time within a trial. To examine the impact of trial length, we added the current number of samples drawn as both an independent predictor and as a interaction with choice type into the above linear regression (Eq. S4). Adding sample number into the model as both a predictor and as an interaction term revealed a significant positive interaction between the decision to stop or continue and how many samples had already been accumulated. Specifically, the decision to continue sampling got faster as the number of samples increased. However, the decision to stop sampling showed the opposite relationship, where the more samples a participant had collected in a trial already, the slower the decision to terminate the trial became (Supplementary Figure 3b).

$$RT = \beta_1(choice\ type) * \beta_2(samples\ drawn) + (1|subj) \qquad \text{(S4)}$$

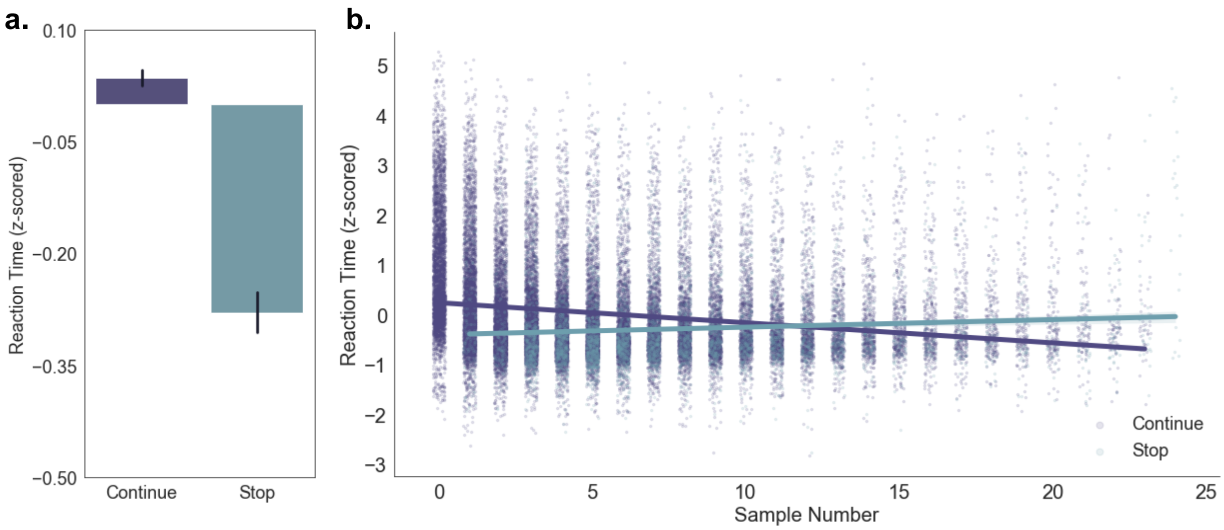

Supplementary Figure 2. Reaction times differ as a function of choice type and sample number. A). Average reaction times (z-scored) aggregated across participants split by choice type (decision to stop sampling or continue sampling). Across participants, the decision to stop sampling took less time (faster RTs) than the decision to continue sampling. B). Choice types share opposing relationships with sample number. The decision to continue gets faster as the number of samples gets larger while the decision to stop gets slower as the number of samples gets larger. Each dot represents a single choice and the color of the dot reflects the choice type (purple = continue sampling, blue = stop sampling). Best fit lines represent slopes for the relationship between reaction times and sample number separated by choice type.
